# Hydrogen-deuterium exchange mass spectrometry of Mtr4 with diverse RNAs reveals substrate-dependent dynamics and interfaces in the arch

**DOI:** 10.1101/2021.09.01.458570

**Authors:** Naifu Zhang, Keith J Olsen, Darby Ball, Sean J Johnson, Sheena D’Arcy

## Abstract

Mtr4 is a eukaryotic RNA helicase required for RNA decay by the nuclear exosome. Previous studies have shown how RNA enroute to the exosome threads through the highly conserved helicase core of Mtr4. Mtr4 also contains an arch domain, although details of potential interactions between the arch and RNA have been elusive. To understand the interaction of *Saccharomyces cerevisiae* Mtr4 with various RNAs, we have characterized RNA binding in solution using hydrogen-deuterium exchange mass spectrometry, and affinity and unwinding assays. We have identified RNA interactions within the helicase core that are consistent with existing structures and do not vary between tRNA, single-stranded RNA, and double-stranded RNA constructs. We have also identified novel RNA interactions with a region of the arch known as the fist or KOW. These interactions are important for RNA unwinding and vary in strength depending on RNA structure and length. They account for Mtr4 discrimination between different RNAs. These interactions further drive Mtr4 to adopt a closed conformation characterized by reduced dynamics of the arch arm and intra-domain contacts between the fist and helicase core.

## Introduction

Eukaryotic RNA processing and surveillance pathways utilize multiple protein complexes to identify and eventually degrade targeted RNA substrates (Chlebowski et al., 2013; Januszyk & Lima, 2014; Schmid & Jensen, 2019; Schneider & Tollervey, 2013). Disruption of these pathways can have deleterious effects on a cell and has been associated with a variety of disease states (Fabre & Badens, 2014; Laffleur & Basu, 2019; Staals & Pruijn, 2010). Mtr4 plays a central role in nuclear RNA processing and surveillance, providing a bridge between substrate targeting and RNA decay (Olsen & Johnson, 2021; E. M. Weick & Lima, 2021). Substrate recognition is performed by Mtr4 directly, and in association with adaptor complexes such as TRAMP, NEXT or PAXT (Schmid & Jensen, 2019). RNA substrates are delivered through Mtr4 to the exosome, which degrades the RNA with 3’ exonuclease and endonuclease activity. Mtr4 prefers RNA substrates with a 4-6 nucleotide, 3’ single-stranded overhang, and accepts a wide range of RNA substrates including rRNA, tRNA, snoRNA, cryptic unstable transcripts, promoter upstream transcripts, and others (Bernstein et al., 2010; Delan-Forino et al., 2017; Jia et al., 2012). The molecular details of how Mtr4 engages with this diverse array of RNAs are poorly understood.

Mtr4 is an essential Ski2-like 3’ to 5’ ATP-dependent RNA helicase (Bernstein et al., 2008; Liang et al., 1996; X. Wang et al., 2008). The protein is composed of a largely unstructured N-terminal tail and β-sheet, and five structured domains (Jackson et al., 2010; Weir et al., 2010). Four of the domains (recA1, recA2, winged-helix, and helical bundle) form a ring-like helicase core (**Figure S1a**). The fifth domain, the arch, is a large insertion within the winged-helix composed of two antiparallel coiled coils (the arm) and a β-barrel (the fist or Kyrpides–Ouzounis–Woese (KOW) motif). The arch is unique to Mtr4 and Ski2, the Mtr4 counterpart in the cytoplasm. The arch extends outward from the helicase core and can adopt various orientations. Comparison of Mtr4 structures, as well as consideration of cryo-electron microscopy (cryo-EM) classes, suggests that the arch conformation is highly dynamic and may be influenced by protein-protein interactions (Jackson et al., 2010; J. Wang et al., 2019; E.-M. Weick et al., 2018; Weir et al., 2010).

RNA binding in the Mtr4 helicase core is primarily mediated by recA1, recA2, and the HB (Schuller et al., 2018; E.-M. Weick et al., 2018; Weir et al., 2010). Protein-RNA interactions involve conserved motifs at the interface of the recA1 and recA2 domains including motifs Ia, Ib, Ic, IV, IVa, and V (**Figure S1a**) (Jankowsky & Fairman-Williams, 2010). RNA binding by these motifs occurs in conjunction with binding and hydrolysis of ATP by motifs Q, I, II, III, Va, and VI. Motif II includes the catalytic aspartic acid and glutamic acid residues. Other notable features are a β- hairpin in recA2 that appears to act as a wedge between RNA strands, as well as a ratchet helix in the helical bundle involved in RNA sequence recognition (Taylor et al., 2014). RNA interactions with the arch are less well defined. The fist in isolation has been shown to interact with structured RNAs, such as dsRNA and tRNA_i_^Met^, but not with ssRNA (Falk et al., 2017; Halbach et al., 2012; Johnson & Jackson, 2013; Weir et al., 2010). The only structural support for arch engagement with RNA in the context of full-length Mtr4 is a low resolution (8.2 Å) cryo-EM map derived from a sub-population of particles where the fist appears to engage with an RNA-DNA substrate simultaneously engaged in the helicase core (E.-M. Weick et al., 2018).

To further investigate *Saccharomyces cerevisiae* Mtr4 binding to various RNA substrates, we have performed hydrogen-deuterium exchange mass spectrometry (HDX), as well as affinity and unwinding assays. We show that RNA interactions in the helicase core are identical between tRNA, single-stranded RNA (ssRNA), and short and long double-stranded RNA (dsRNA) substrates. These interactions are consistent with previous structural studies, but do not account for different affinities to Mtr4. However, we also detect interactions between the fist and RNA that do vary depending on RNA structure and length. These interactions reduce the dynamics of the arm and place Mtr4 in a closed conformation with the fist near the surface of recA2. We show that removal of the fist compromises both RNA binding and unwinding. Taken together, our data reveals an important role for the arch in RNA recognition and demonstrates a connection between arch-RNA interactions and unwinding activity.

## Results

To characterize the solution interaction between *S. cerevisiae* Mtr4 and RNA we employed HDX. We recovered over 300 peptides redundantly spanning 81% of the Mtr4 sequence (**Tables S1-S2**). The longest stretch lacking peptides was the disordered N-terminal tail (**Figure S1c**). We monitored the deuterium uptake of all peptides in Mtr4 alone and bound by RNA. For Mtr4 alone, the absolute amount of deuterium uptake correlated well with known secondary and tertiary structural elements. Deuterium uptake was lowest in the globular recA domains and highest in the N-terminal sheet and parts of the arch. The N-terminal sheet and arch are likely mobile and solvent exposed when Mtr4 is alone in solution. This is in keeping with the various orientations of the arch observed in structures of Mtr4 from yeast, human and plants (Conrad et al., 2016; J. Wang et al., 2019; Weir et al., 2010).

### tRNA_i_^Met^ binds Mtr4 at the interface of the recA1 and recA2 domains

To identify changes in Mtr4 upon binding RNA, we first compared deuterium uptake of Mtr4 alone to that of Mtr4 with a hypomodified tRNA_i_^Met^. Hypomodified tRNA_i_^Met^ is a native substrate of Mtr4 and our construct had a 12-nucleotide, 3’ overhang with sequence A_9_CGC (**Table S3**) (X. Wang et al., 2008). We determined the dissociation constant (K_d_) between full-length Mtr4 (Mtr4^WT^) and tRNA_i_^Met^ to be 0.27 ± 0.02 µM (± SD; **Table 1**). Based on this we used a 5.4-fold excess of tRNA_i_^Met^ to ensure at least 90% of Mtr4^WT^ was RNA-bound. Throughout this work, we show the difference in deuterium uptake between Mtr4 alone and RNA-bound using heatmaps created with HD-eXplosion (Zhang et al., 2021). The red shading indicates a statistically significantdifference or protection upon RNA binding. We defined significance as a decrease ≥0.5 Da and a p-value ≤00.01 in a Welch’s t-test (n=3). Such significant decreases in deuterium uptake upon addition of tRNA_i_^Met^ were observed in all domains of Mtr4^WT^, except the N-terminal sheet and the winged-helix (**Figure 1a, d**).

**Table 1.**
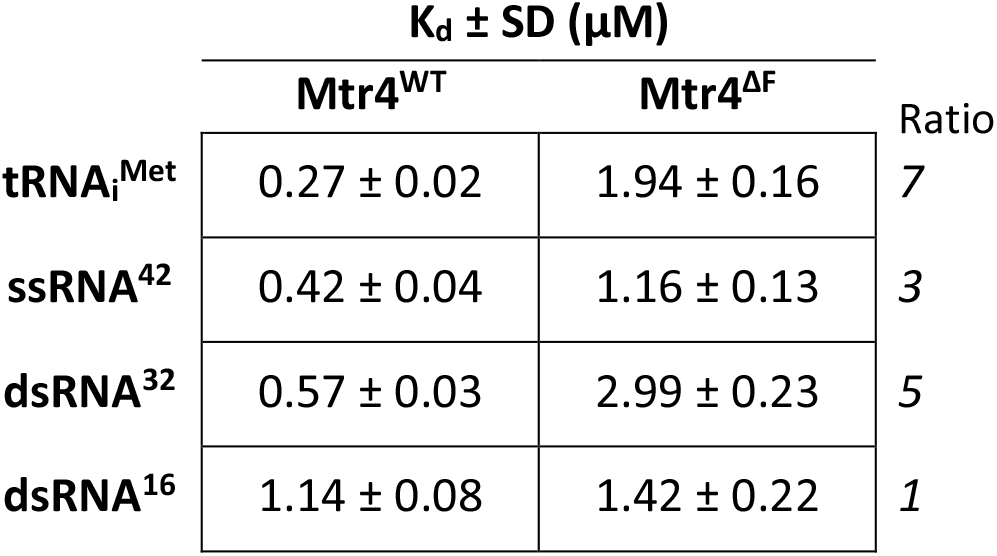
Dissociation constants of Mtr4^WT^ or Mtr4^ΔF^ to various RNAs. The dissociation constants (K_d_) between Mtr4^WT^ or Mtr4^ΔF^ and tRNA_i_^Met^, ssRNA^42^, dsRNA^32^, or dsRNA^16^ were measured using an electrophoretic mobility shift assay. K_d_ values are an average of three independent gels plus or minus one standard deviation (SD). The ratio between Mtr4^ΔF^ and Mtr4^WT^ is listed to one significant figure.

**Figure 1.**
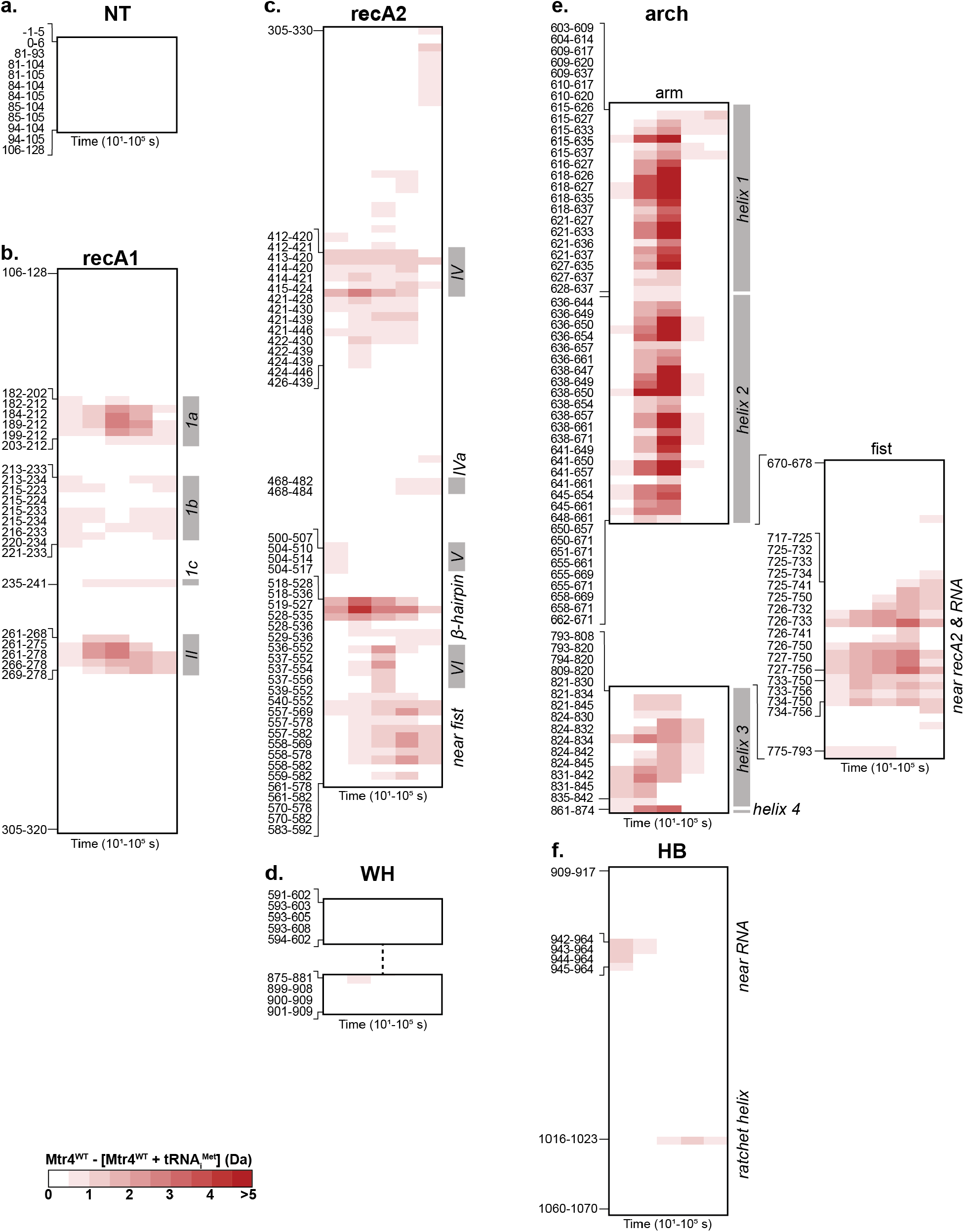
Addition of tRNA_i_^Met^ caused reduced deuterium uptake in several domains of Mtr4^WT^. Heatmaps showing the difference in deuterium uptake between Mtr4^WT^ alone and Mtr4^WT^ with 5.4-foldtRNA_i_^Met^. Peptides were recovered from the following domains: (**a**) N-terminal (NT), (**b**) recA1, (**c**) recA2,(**d**) winged-helix (WH), (**e**) arch, and (**f**) helical bundle (HB). The time points of exchange (x-axis) were 10 s, 10^2^ s, 10^3^ s, 10^4^ s, and 10^5^ s. Red blocks indicate a difference ≥0.5 Da with a p-value ≤0.01 in a Welch’s t-test (n=3). Boundaries of peptides with significant changes are shown (left y-axis) and Mtr4 features are labeled (right y-axis). Example uptake plots and a full list of peptides are in **Figure S1b-c**. Figure was created using HD-eXplosion (Zhang et al., 2021). In this study, Mtr4 numbering does not include the start methionine (**Table S2**).

The decreases in deuterium uptake seen in the helicase core upon addition of tRNA_i_^Met^ are consistent with previous RNA-bound Mtr4 structures. The ‘ core’ refers to the domains shared with most other helicases and includes the recA1, recA2, winged-helix and helical bundle domains (**Figure S1a**). In the recA1, the largest protection occurred in two regions near the RNA (**Figure 1b, 2**). The first region contains motif 1a directly involved in RNA binding, while the second region is near motif II and the catalytic residues. Protection was also seen for RNA-binding motifs 1b and 1c, albeit less than motif 1a. In the recA2, protection similarly occurred in motif IV situated near the RNA backbone, and a β-hairpin wedge that purportedly acts to split RNA strands (**Figure 1c, 2**) (Büttner et al., 2007; E.-M. Weick et al., 2018). Protection was also seen for motifs IVa, V and VI, but less than motif IV and the β-hairpin wedge. Small decreases also occurred in the helical bundle, specifically in a region that sits atop the RNA bases, and in the single peptide (1016-1023) near the ratchet helix that recognizes the 3’ overhang (**Figure 1f, 2**) (Taylor et al., 2014).

The broad agreement between high-resolution structures and the regions protected from deuterium exchange reinforces that the tRNA_i_^Met^ 3’ overhang binds at the interface of the recA1 and recA2 (**Figure 2**). The data show that this occurs in solution and in the context of a large and folded tRNA substrate for which a co-structure with Mtr4 is not available. Motifs 1a, II, IV, and the β-hairpin wedge emerge as being the most important for RNA binding, as within the helicase core they show the largest protection at any one time point (**Figure 1b-d, f**). These regions are also characterized by a particular kinetic profile with protection being sustained throughout almost the entire HDX time course. The sustained decrease is highlighted by uptake plots of example peptides (**Figure S1b**). Such a kinetic profile is thus likely characteristic of an RNA-binding region in Mtr4.

**Figure 2.**
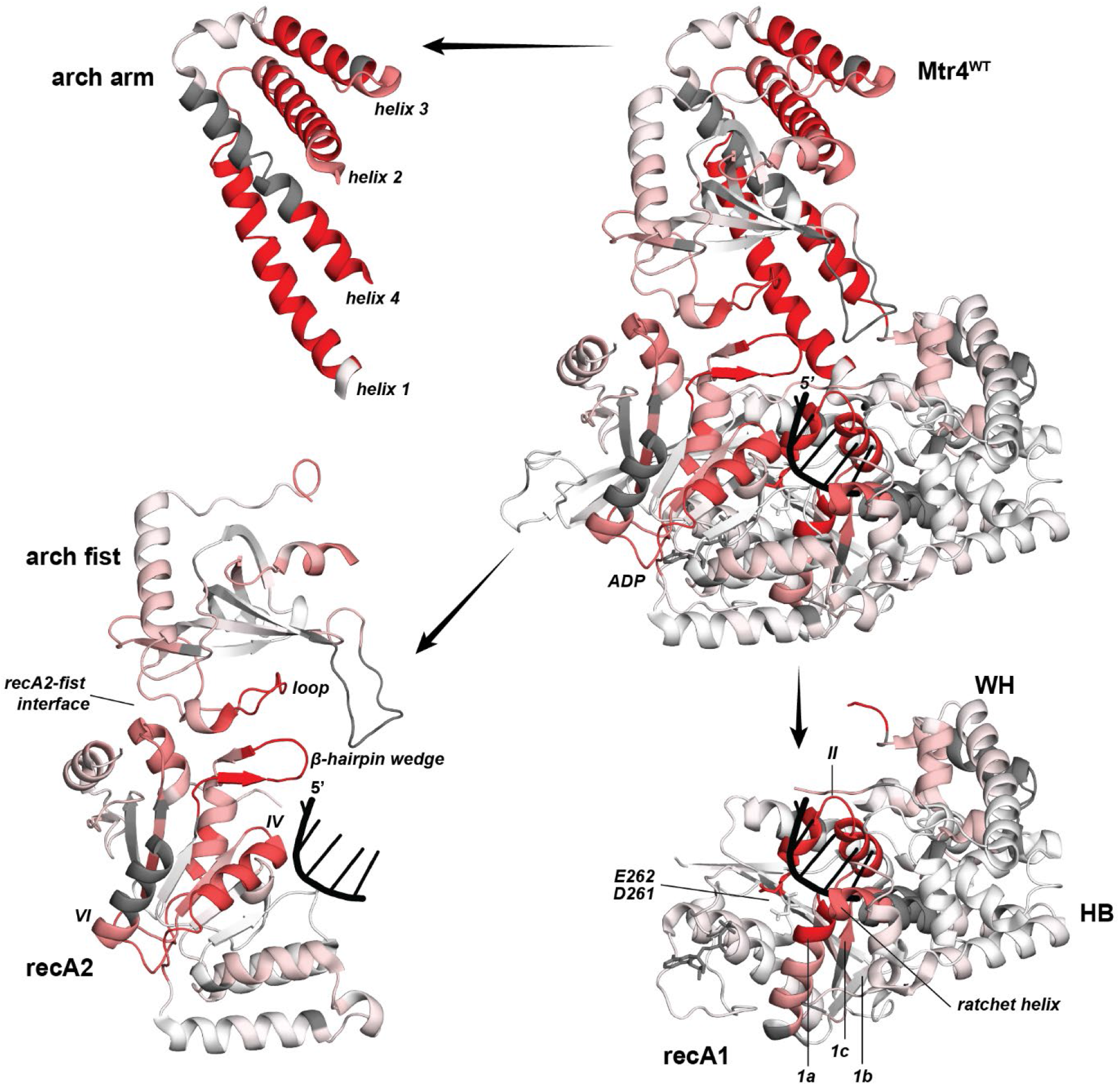
The helicase core and arch of Mtr4^WT^ showed reduced deuterium uptake in the presence of tRNA_i_^Met^. Mtr4 (PDB 2XGJ Chain A) is colored by the difference in fractional deuterium uptake between Mtr4^WT^ alone and Mtr4^WT^ with 5.4-fold tRNA_i_^Met^ after 10^3^ s exchange. The scale is 0-20% (white to red) and is based on DynamX residue-level scripts without statistical filtering from the data shown in **Figure 1**. Residues without coverage are grey and Mtr4^WT^ features are labeled including ADP (gray sticks), RNA (black), and D261-E262 (sticks). The images depict the entire Mtr4^WT^ structure (top right), the arm of the arch (top left), the fist and recA2 (bottom left), and the recA1, WH, and HB (bottom right).

### Binding of tRNA_i_^Met^ stabilizes a closed conformation of Mtr4

Addition of tRNA_i_^Met^ to Mtr4^WT^ also caused a decrease in deuterium uptake in the arch. The arch contains an arm composed of two sets of anti-parallel helices (designated here as helices 1 and 4; and helices 2 and 3) with a fist domain inserted between helices 2 and 3 (**Figure S1a**). Structural work has shown that the arch can adopt various positions relative to the core. In some Mtr4 structures the fist of the arch is positioned near to recA2 in a closed conformation, while in others the fist is solvent exposed to varying degrees (Olsen & Johnson, 2021). Upon addition of tRNA_i_^Met^, we observed a decrease in uptake in both the fist and the arm of the arch (**Figure 1e**).Within the fist, the largest protection occurred in a loop that in closed Mtr4 forms a continuous surface with the established RNA-binding site and β-hairpin wedge (**Figure 2**). This protection is likely attributed to direct RNA-binding due to the loop’s location, as well as the kinetic profile of protection. Protection of the loop persisted throughout the entire HDX time course, similar to the known RNA-binding regions (**Figure 1e, S1b**). Direct binding of the fist to dsRNA has previously been reported using an isolated fist construct (Falk et al., 2017). Reduced deuterium uptake also extended to the opposite side of the fist, distal to the RNA (**Figure 2**). In closed Mtr4 this opposite side interacts with a surface of recA2, and we observed a similar decrease in uptake at this recA2 surface. This suggests that the fist contacts the recA2 in the presence of tRNA_i_^Met^ (**Figure 1c, e**). It seems likely that the fist directly binds the tRNA_i_^Met^ through its’ loop and is then positioned near recA2, putting Mtr4 in a closed conformation.

The data further show that tRNA_i_^Met^ causes a drastic change in the conformational plasticity of the four helices of the arm. Of all the Mtr4^WT^ regions, the arm had the largest reduction in deuterium uptake upon addition of tRNA_i_^Met^ (**Figure 1e, 2**). The kinetic profile of the decreased uptake was also distinct from that associated with RNA binding by the helicase core and fist. Rather than showing a sustained decrease throughout the entire HDX time course, the arm showed a relatively short-lived decrease that appeared at 10^2^ s and was diminished by 10^4^ s (**Figure 1e, S1b**). This, combined with the fact that previous structures show no direct association between the arm and RNA, suggests the protection to be a result of a conformational change. Binding to the tRNA_i_^Met^ seems to constrain the arch to one or a subset of conformations due to contacts formed between the fist and both tRNA_i_^Met^ and recA2, placing Mtr4 into a closed conformation.

### RNA interactions outside the helicase core account for Mtr4 substrate preference

We next characterized Mtr4 binding to ssRNA. We used a 42-nucleotide ssRNA with a polyadenylated tail at its 3’ end (ssRNA^42^) (**Table S3**). The 3’ poly(A) included five adenines as this is the ideal length to maximize binding to Mtr4 (Jia et al., 2011). ssRNA^42^ is not predicted to have any secondary structure or to dimerize (based on IDT OligoAnalyzer tool) and was designed to be long enough to traverse the helicase core and the fist. We determined the affinity of ssRNA^42^ to Mtr4^WT^ to be 0.42 ± 0.04 μM (± SD; **Table 1**); 1.6-fold weaker than the binding of tRNA_i_^Met^ (**Table 1**). This reduced binding, combined with large amounts of ssRNA^42^ being problematic in our HDX system, limited our ability to saturate Mtr4^WT^ during the exchange reaction. Instead, we used a 2-fold excess of ssRNA^42^ over Mtr4^WT^ and repeated the experiment with tRNA_i_^Met^ under these same conditions (all peptides shown in **Figure S2**). The dataset had less usable peptides than the previous dataset due to RNA-induced peptide carryover but still contained almost 200 peptides covering 58% of the Mtr4^WT^ sequence (**Table S1-S2**). This facilitated direct comparison of Mtr4^WT^ binding to ssRNA^42^ and tRNA_i_^Met^. The Mtr4^WT^ regions altered by tRNA_i_^Met^ were broadly similar when it was 2-fold and 5.4-fold (compare **Figure 1 and S2**; **Figure 6c**).

Comparison of ssRNA^42^ and tRNA_i_^Met^ revealed identical decreases in deuterium uptake in the core RNA-binding elements of Mtr4^WT^ (**Figure 3, black**). Important motifs 1a, IV, the β-hairpin wedge, and the ratchet helix, all showed the same degree of protection upon addition of either ssRNA^42^ or tRNA_i_^Met^. While the involvement of the same regions is not surprising, the identical magnitude of protection was unexpected given the difference in their binding affinities for Mtr4^WT^ (**Table 1**). This demonstrates that the difference must be due to binding interactions outside of the Mtr4^WT^ helicase core. Outside of the helicase core, addition of ssRNA^42^ caused less protection than tRNA_i_^Met^ (**Figure 3, gray**). Less protection was seen in the fist and arm of the arch, as well as at the surface of recA2. The smaller effect of ssRNA^42^ compared to tRNA_i_^Met^ outside the helicase core correlates with their relative affinities for Mtr4^WT^. Thus, while tight RNA-binding by Mtr4 primarily comes from interactions in the helicase core, regions outside the core clearly play a role in the fine tuning of Mtr4 substrate preference.

**Figure 3.**
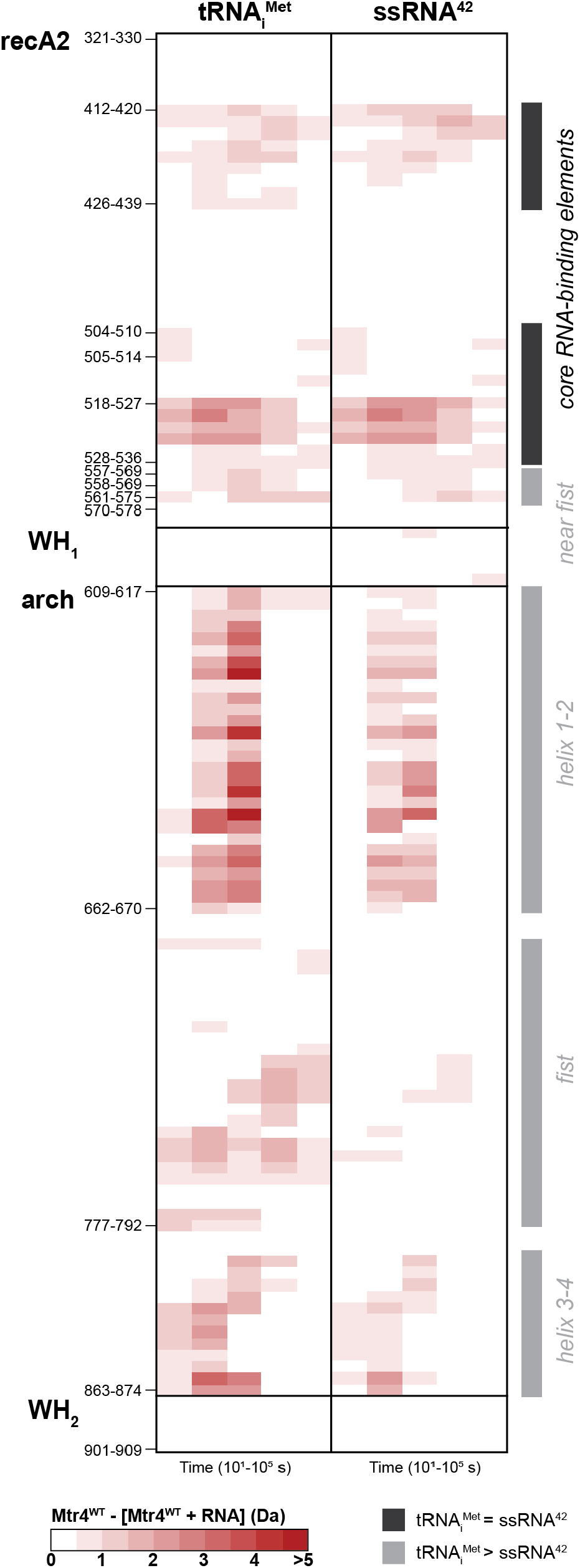
Protection in the arch of Mtr4^WT^ was reduced for ssRNA^42^ compared to tRNA_i_^Met^. Heatmaps showing the difference in deuterium uptake between Mtr4^WT^ alone and Mtr4^WT^ with 2-fold tRN^iMet^ or ssRNA^42^. Peptides from the recA2, WH (split into WH_1_ and WH_2_), and arch are shown with domain boundaries and key peptides (left y-axis) indicated. The complete data set is shown in **Figure S2**. The time points of exchange (x-axis) were 10 s, 10^2^ s, 10^3^ s, 10^4^ s, and 10^5^ s. Red blocks indicate a difference ≥0.5 Da with a p-value ≤0.01 in a Welch’s t-test (n=3). Mtr4 features are labeled (right y-axis) with black indicating equal difference between tRNA_i_^Met^ and ssRNA^42^ and gray indicating less difference for ssRNA^42^ than tRNA_i_^Met^. Figure was created using HD-eXplosion (Zhang et al., 2021).

The data with ssRNA^42^ provides several additional insights into RNA binding by Mtr4^WT^. The first is that the fist does in fact form contacts with ssRNA^42^ as the ssRNA^42^ did induced protection, albeit less than tRNA_i_^Met^ (**Figure 3**). This builds on previous work showing that the isolated fist can bind dsRNA, but not ssRNA (Halbach et al., 2012; Johnson & Jackson, 2013). Contacts between the fist and ssRNA are likely promoted in full-length Mtr4 as the RNA would be tethered nearby by tight interactions in the helicase core. Another insight is the apparent connection between the arm and fist of the arch, and the surface of recA2. These regions showed a very similar reduction in protection when tRNA_i_^Met^ was replaced with ssRNA^42^ (**Figure 3**). This suggests that RNA-binding by the fist, conformational change in the arm, and interactions between the fist and recA2 surface, are coupled events that result in RNA-bound Mtr4 adopting a closed conformation. Further, this closed conformation appears grossly similar for Mtr4 bound by either tRNA_i_^Met^ or ssRNA^42^. The latter is indicated by the similar pattern of protection upon addition of either type of RNA (**Figure 3**).

### The Mtr4 arch senses RNA length and structure

To further characterize the RNA features recognized by Mtr4, we made use of two dsRNA constructs. These constructs have been used to characterize the nucleic acid unwinding rate by Mtr4 (**Table 2**) (Jia et al., 2011). One construct had a 16-bp duplex with a 6-nucleotide, 3’ overhang of UAAAAA, which is approximately the same length as the acceptor stem, T-arm, and T-loop of the tRNA_i_^Met^ (dsRNA^16^; **Table S3**). The other construct was similar but had an extended duplex (32 bp) with a hairpin and a nick (dsRNA^32^; **Table S3**). The dsRNA^32^ is approximately the same number of base pairs as tRNA_i_^Met^, although it likely adopts a different fold. Both dsRNA^32^ and dsRNA^16^ had reduced binding to Mtr4^WT^ compared to tRNA_i_^Met^. The affinities were 0.57 ±0.03 and 1.14 ± 0.08 μM (± SD; **Table 1**); 2-fold and 4-fold weaker than tRNA_i_^Met^ respectively. Reduced binding of dsRNA^32^ compared to tRNA_i_^Met^ shows that Mtr4 affinity is influenced by the fold of the RNA, while reduced binding of dsRNA^16^ compared to dsRNA^32^ shows that dsRNA length is also an important determinant of binding.Given the different affinities of the dsRNA constructs, we performed another HDX experiment to directly compare their binding to Mtr4^WT^. We again used a 2-fold excess of dsRNA over Mtr4^WT^ and included both tRNA_i_^Met^ and ssRNA^42^ for direct comparison (**Table S1-S2;** all peptides shown in **Figure S3a-b**). The Mtr4^WT^ regions altered by tRNA_i_^Met^ and ssRNA^42^ were similar between replicate experiments (compare **Figure S2 and S3a-b**; **Figure 6c**). Concurrent analysis of all four RNA constructs showed identical decreases in deuterium uptake in the core RNA-binding elements of Mtr4^WT^ (**Figure S3a-b**). RNA-binding motifs 1a, 1c, IV, and the ratchet helix, all had the same degree of protection regardless of the type of RNA added (example peptides shown in **Figure 4**). This shows that dsRNA^16^, dsRNA^32^, ssRNA^42^, and tRNA_i_^Met^ all bind to the helicase core in the same manner and with the same affinity. RNA features such as fold and length, seem to have little to no effect on Mtr4 binding to the 3’ RNA tail. This reinforces the conclusion that differences in RNA affinity for Mtr4^WT^ are dictated by interactions outside of the helicase core.

**Table 2.** Unwinding rate of Mtr4^WT^ or Mtr4^ΔF^ with dsRNA^16^ or dsRNA^32^ substrates. The maximum strand-separation rate constants (k_uwn_^max^) for Mtr4^WT^ or Mtr4^ΔF^ were measured using dsRNA^32^ or dsRNA^16^ as substrate. k_uwn_^max^ values are an average of three independent experiments plus or minus one standard deviation (SD). The ratio between Mtr4^ΔF^ and Mtr4^WT^ is listed to one significant figure. The data for Mtr4^WT^ with dsRNA^16^ is from previously published work (Taylor et al., 2014).

| | $k_{\text{uwn}}^{\text{max}} \pm \text{SD} (\text{min}^{-1})$ | | Ratio |
| --- | --- | --- | --- |
|  | Mtr4<sup>WT</sup> | Mtr4<sup>ΔF</sup> |  |
| dsRNA<sup>32</sup> | 0.56 ± 0.04 | 0.25 ± 0.02 | 0.44 |
| dsRNA<sup>16</sup> | 0.59 ± 0.05<sup>a</sup> | 0.56 ± 0.07 | 0.95 |

<sup>a</sup>Previously published (Taylor et al., 2014)

**Figure 4.**
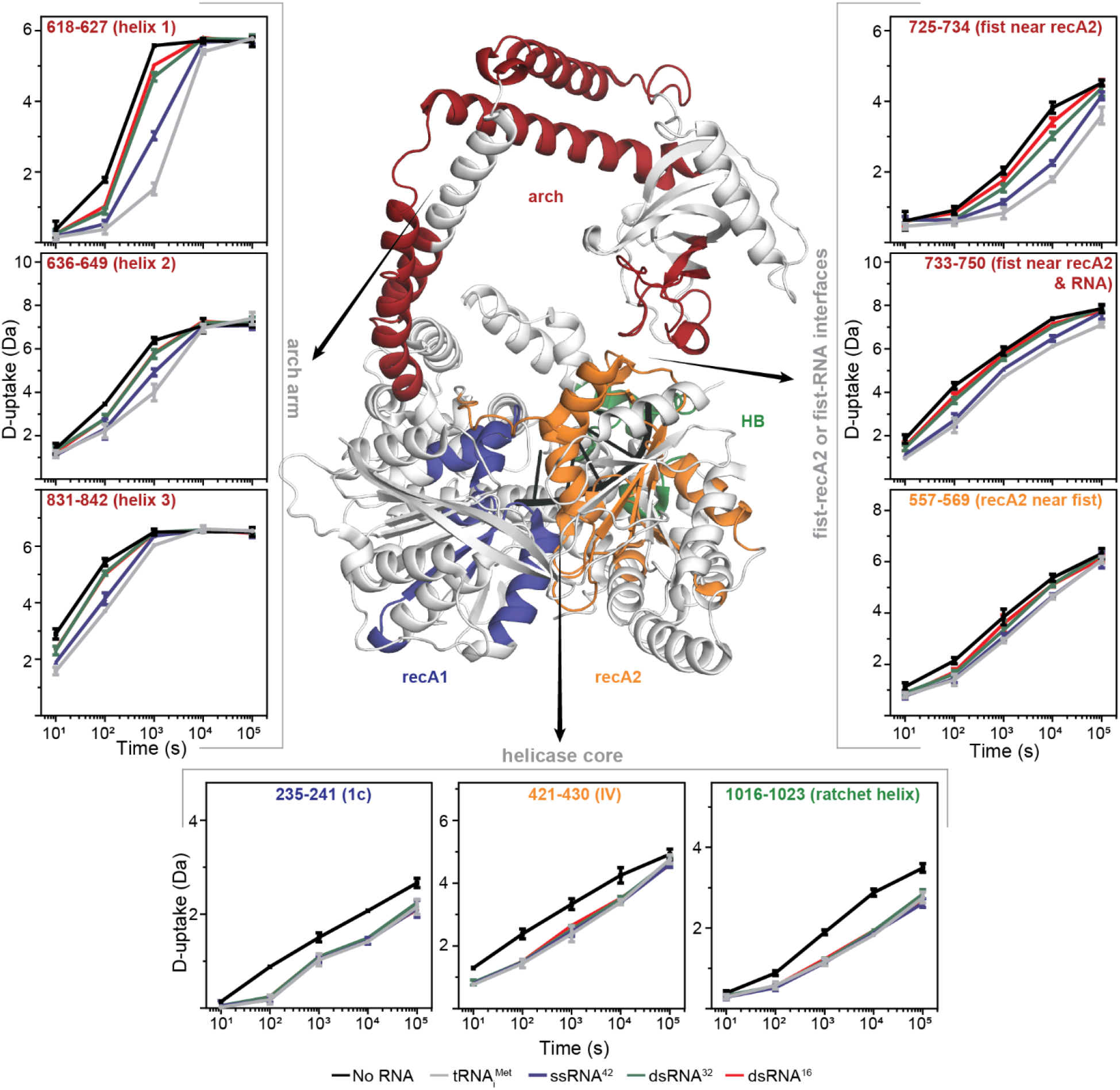
Deuterium uptake in the arch, but not the helicase core, of Mtr4^WT^ was influenced by RNA structure and length. Mtr4 (PDB 2XGJ Chain A) surrounded by example uptake plots from the arch arm (left), fist and recA2 surface (right), and helicase core (bottom). Uptake plots show Mtr4^WT^ alone (black) and with 2-fold tRNA_i_^Met^ (gray), ssRNA^42^ (blue), dsRNA^32^ (green), or dsRNA^16^ (red). Plotted uptake is the average of three replicates and error bars are plus or minus two standard deviations. The y-axis range is 80% of theoretical maximum uptake, assuming the N-terminal residue undergoes complete back-exchange. Data have not been corrected for back-exchange. Heatmap for all peptides from this dataset is in **Figure S3**. Colors on the Mtr4 structure show regions protected by 5.4-fold tRNA_i_^Met^ (based on data in **Figure 1**) in the recA1 (blue), recA2 (orange), arch (red), and HB (green) domains.

In contrast to the helicase core, the degree of protection in the arch was less for both dsRNAs compared to either the ssRNA^42^ or tRNA_i_^Met^. This reduced protection was seen in all helices of the arm, the fist, and at the surface of recA2 (**Figure 4, 5, and S3a-b**). The reduced protection is in line with the weaker binding affinities of the dsRNAs (**Table 1**) and suggests that the fist has less, or more transient, interactions with the dsRNAs compared to ssRNA^42^ or tRNA_i_^Met^. The reduced protection in the arm, fist, and recA2 surface reinforces the connection between RNA-binding and arch movement that favors a closed conformation of Mtr4. Close inspection of the data further shows greater protection for dsRNA^32^ than dsRNA^16^ (**Figure 5**). This is clearer when a cutoff of 0.3 Da (rather than 0.5 Da) is used (**Figure S3c**). A cutoff of 0.3 Da is justified as the standard error of the mean with a student’s t-distribution value for 0.05 significance, gives approximately 0.22 Da for each of the data sets (Hageman & Weis, 2019). Altogether, comparison of the four RNA substrates shows unequivocally that the contribution of the arch depends on both RNA length and structure.

**Figure 5.**
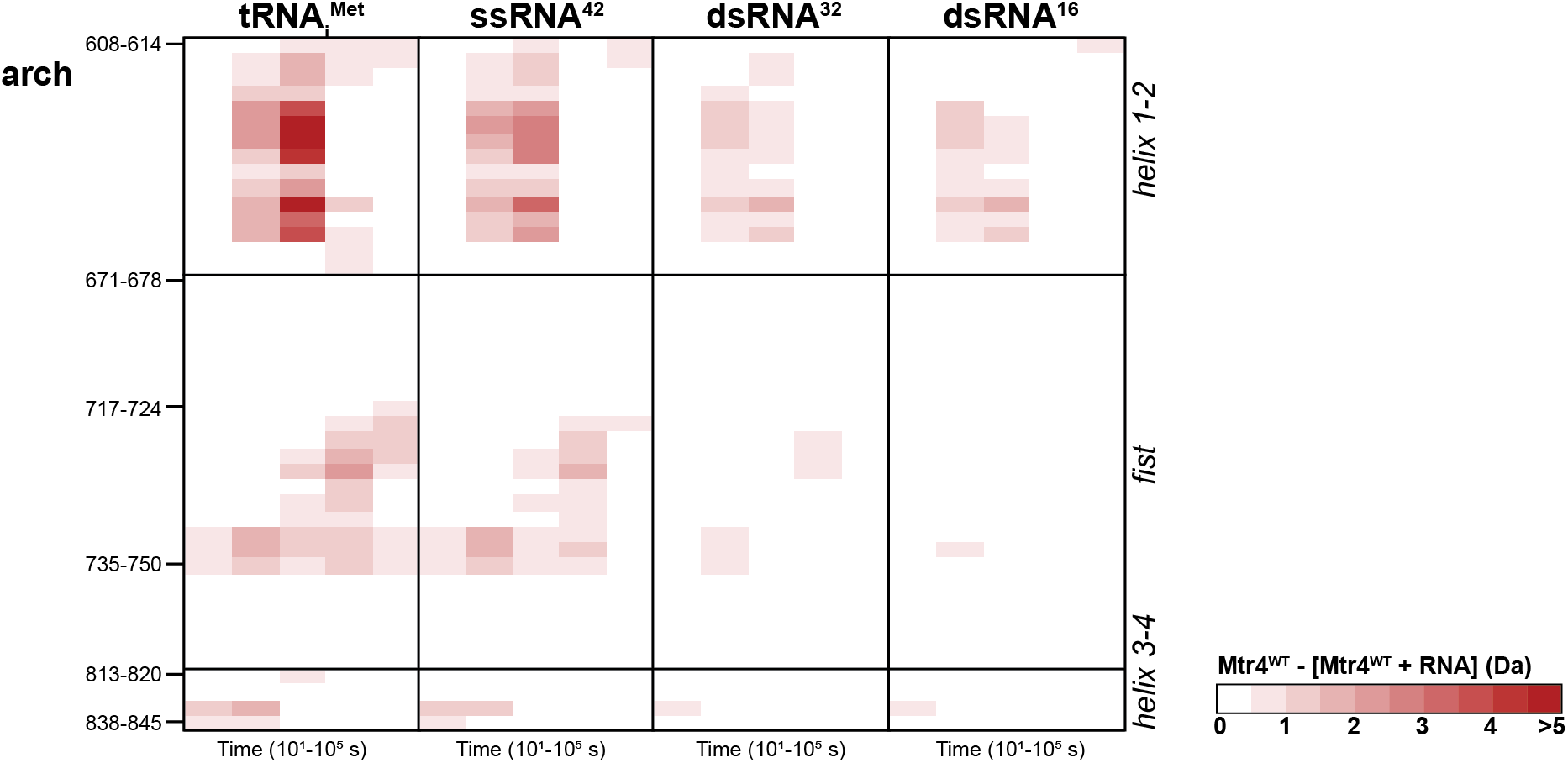
The degree of protection in the arch of Mtr4^WT^ varied with RNA type. Heatmaps showing the difference in deuterium uptake between Mtr4^WT^ alone and Mtr4^WT^ with 2-fold tRNA_i_^Met^, ssRNA^42^, dsRNA^32^, or dsRNA^16^, for peptides from the arch. The complete data set is shown in **Figure S3**. The time points of exchange (x-axis) were 10 s, 10^2^ s, 10^3^ s, 10^4^ s, and 10^5^ s. Key peptides (left y-axis) and Mtr4 features (right y-axis) are labeled. Red blocks indicate a difference ≥0.5 Da with a p-value ≤0.01 in a Welch’s t-test (n=3). Figure was created using HD-eXplosion (Zhang et al., 2021).

### Fist interactions with RNA drive Mtr4 conformational change

In parallel to varying the RNA substrate, we characterized a fist-less Mtr4 (Mtr4^ΔF^) with residues 677-821 replaced by a three-glycine linker. Removal of the fist reduced binding of tRNA_i_^Met^, ssRNA^42^, and dsRNA^32^ by 7-, 3-, and 5-fold respectively (**Table 1**). This reduction suggests that the fist directly contributes to recognition of these RNAs and is consistent with the similar kinetics of protection seen for the helicase core and the fist in the HDX with Mtr4^WT^ (**Figure 1 and 3**). The larger effect of fist removal on binding dsRNA^32^ compared to ssRNA^42^ was surprising and may be due to binding differences not apparent when using Mtr4^WT^. Unlike tRNA_i_^Met^, ssRNA^42^ and dsRNA^32^, the removal of the fist did not significantly affect Mtr4 binding to dsRNA^16^. This shows that the fist plays little to no role in recognition of dsRNA^16^ and is keeping with the low levels of protection seen for the fist in the HDX with Mtr4^WT^ (**Figure 5**). The 1.5-2.0 μM affinities for Mtr4^ΔF^likely represent RNA-binding in the helicase core, with the tighter affinities of Mtr4^WT^ being attributable to RNA interactions with the fist (**Table 1**). The affinity differences between Mtr4^ΔF^ and Mtr4^WT^ thus demonstrate how the fist is directly responsible for fine tuning binding preference depending on RNA features.

We also characterized Mtr4^ΔF^ binding to the four different RNAs using HDX in the same conditions as used for Mtr4^WT^. In fact, Mtr4^ΔF^ and Mtr4^WT^ were exchanged and quenched simultaneously, with the Mtr4^ΔF^ samples being thawed and injected into the mass spectrometer immediately following the Mtr4^WT^ samples (**Table S1-S2;** all peptides shown in **Figure S4a-b**). When compared to Mtr4^WT^ in the absence of RNA, Mtr4^ΔF^ had similar deuterium uptake in the helicase core and increased uptake in the arch. Compare, for example, peptide 235-241 from the core and peptide 636-649 from the arch, between Mtr4^WT^ (**Figure 4**, black trace) and Mtr4^ΔF^ (**Figure 6b**, black trace). This shows that deletion of the fist did not compromise the integrity of the enzymatic core but did increase the dynamics of the arch only when Mtr4 was alone in solution.

The changes in deuterium uptake of Mtr4^ΔF^ upon addition of RNA were identical for all four of our RNA constructs, but differences were observed between Mtr4^ΔF^ and Mtr4^WT^. Reduced uptake was seen in the helicase core of Mtr4^ΔF^, with equal protection seen upon addition of tRNA_i_^Met^, ssRNA^42^, dsRNA^32^, or dsRNA^16^ (**Figure 6b and S4a-b**). While coverage of some RNA-binding motifs was lacking, protection was seen for motifs 1a, 1c, IV, and the ratchet helix (**Figure 6c**). For these motifs, particularly 1a, the degree of protection was slightly less for Mtr4^ΔF^ than Mtr4^WT^ (**Figure 6a**, compare left and right). The interactions between the fist and RNA may thus subtly alter the stability and trajectory of RNA through the helicase core. A more obvious difference was in the arm, as the large RNA-induced protection seen in Mtr4^WT^ was completely lost in Mtr4^ΔF^. All helices of the arm and the surface of the recA2 of Mtr4^ΔF^ had no change in uptake upon addition of RNA (**Figure 6**). Removal of the fist clearly prevents the conformational change of the arch revealing that RNA interactions with the fist and not the helicase core drive arch movements. Comparison of Mtr4^WT^ and Mtr4^ΔF^ identifies a clear role for the fist in binding RNA and regulating Mtr4 transition between an open, unbound state and a more-closed, bound state.

**Figure 6.**
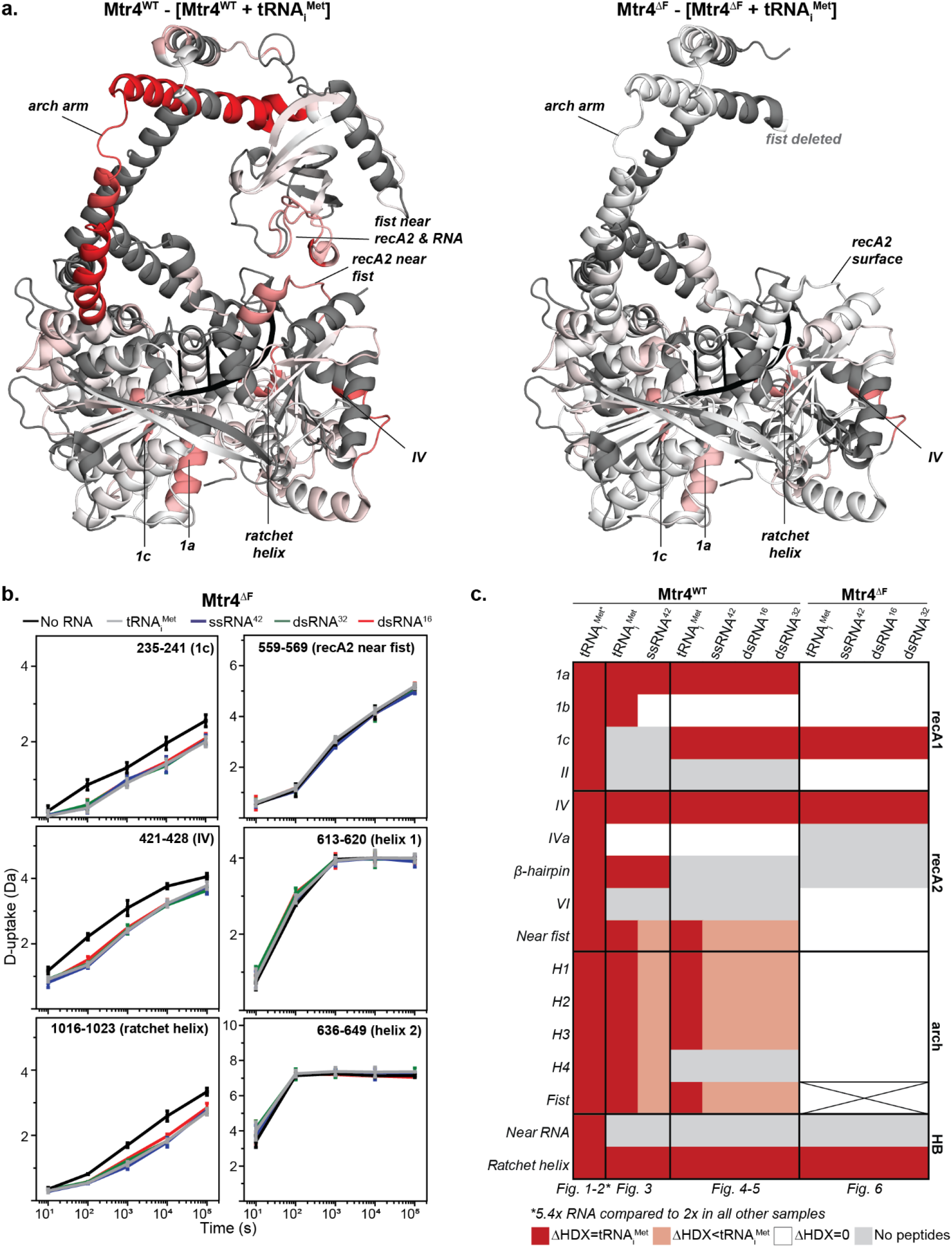
Deletion of the fist in Mtr4^ΔF^ caused a loss of protection in the arch. **(a)** Mtr4 (PDB 2XGJ Chain A) is colored by the difference in fractional deuterium uptake between Mtr4^WT^ (left) or Mtr4^ΔF^ (right) alone and with 2-fold tRNA_i_^Met^ after 10^3^ s exchange. The scale is 0-20% (white to red) and is based on DynamX residue-level scripts without statistical filtering. Residues without coverage are gray. Complete data for Mtr4^ΔF^ with each RNA is in **Figure S4. (b)** Example uptake plots with Mtr4^ΔF^ alone (black) and with 2-fold tRNA_i_^Met^ (gray), ssRNA^42^ (blue), dsRNA^32^ (green), or dsRNA^16^ (red). Plotted uptake is the average of three replicates and error bars are plus or minus two standard deviations. The y-axis range is 80% of theoretical maximum uptake, assuming the N-terminal residue undergoes complete back-exchange. Data have not been corrected for back-exchange. **(c)** Summary of all HDX experiments. Mtr4 features (left y-axis) are annotated as having equal (red) or less (light red) protection as the tRNA_i_^Met^ sample in the same HDX experiment, no protection (white), or no coverage (gray). The fist of Mtr4^ΔF^ is crossed.

To fully ascertain the role of the fist, we used pre-steady state assays to measure the unwinding rate of Mtr4^WT^ and Mtr4^ΔF^. We compared dsRNA^32^ and dsRNA^16^ substrates as the fist plays a small and large role in their binding respectively (**Table 1**). Mtr4^WT^ and Mtr4^ΔF^ unwound the dsRNA^16^ at similar speeds with maximum strand-separation constants at enzyme saturation being ∼0.6 min^−1^ (**Table 2**). This is consistent with the fist playing only a minor role for the short dsRNA^16^. A similar result was obtained with a completely arch-less Mtr4 (Jackson et al., 2010). Mtr4^WT^ and Mtr4^ΔF^ however differed in their ability to unwind dsRNA^32^, with the deletion mutant being 2-fold slower (**Table 2, Figure S5**). This shows that the fist contributes to the catalytic unwinding activity that occurs in the helicase core of Mtr4. The interactions between the fist and RNA that drive conformational change of the arch are thus relevant to both the binding and unwinding of RNA by Mtr4.

## Discussion

Mtr4 plays an important gatekeeping role for RNA decay in the nucleus, engaging a wide range of RNAs and delivering unwound substrates to the exosome for degradation (Delan-Forino et al., 2017; E.-M. Weick et al., 2018). Prior to this work, the molecular underpinnings of Mtr4 preference among the many diverse RNA substrates were not clear. Our solution studies, however, demonstrate a mechanism whereby substrate selection is driven by the arch. As most of our experiments were conducted in the absence of ATP, our observations are relevant to initial RNA binding and recognition events.

In our HDX experiments, we detect RNA interactions with the helicase core that are consistent with known Mtr4-RNA structures (**Figure 1 and 2**). The observed binding in the core is identical between all RNA substrates. This is notable as our various RNAs do not have identical 3’ overhangs. The tRNA has an overhang of A_9_CGC, while the other RNAs have A_5_ (**Table S3**). Any differences in how these overhangs bind the core, if any, must be subtle enough to evade detection in our assay. It is also notable that the RNA features beyond the 3’ overhang do not influence binding in the core in an appreciable way. tRNA, for example, binds the core in an identical fashion to a 42-nt ssRNA. Most significantly, however, similar binding in the core demonstrates that determinants of substrate selectivity, as reflected in varied binding affinities (**Table 1**), must occur elsewhere, namely in the arch.

Our HDX experiments also reveal RNA-induced impacts in both the arm and fist of the arch. The magnitude of these impacts varies depends on RNA type; with tRNA having the largest impact, followed by ssRNA, long dsRNA, and lastly, short dsRNA (**Figure 4 and 5**). Little to no changes occur in the arch upon the addition of short dsRNA. This shows that arch involvement is a function of both RNA structure and length. Based on kinetic profiles of exchange, we ascribe the changes in the fist to direct interactions with the RNA. This is consistent with previous work showing that a fist-only construct can interact with RNA (Halbach et al., 2012; Weir et al., 2010). Moreover, the data imply that fist-RNA interactions are enhanced in the context of full-length Mtr4. This is because we observe significant fist interactions with ssRNA (**Figure 3**), whereas no ssRNA interactions were previously observed for the fist-only construct. Presumably, these interactions are enhanced by the tethering of the RNA to the nearby helicase core.

We further show that fist-RNA interactions are relevant to helicase activity. Removal of the fist reduces the ability of Mtr4 to unwind a 32-bp dsRNA that interacts with the fist but has no effect on unwinding a 16-bp dsRNA that does not interact with the fist (**Table 2**). This provides an explanation for an earlier observation that removal of the arch has minimal impact on unwinding the 16-bp substrate (Jackson et al., 2010), and further clarifies the molecular basis of previous conclusions that the arch plays a role in unwinding events (Taylor et al., 2014). Direct RNA interactions with the fist appear to facilitate unwinding across larger or more complex substrates. Since Mtr4 will most likely encounter substrates larger than a 16-bp dsRNA in vivo, we expect that the arch will routinely be involved in unwinding activity. Our model is consistent with a single-molecule study showing that Mtr4 translocates along an RNA strand only when duplex RNA is located immediately upstream (Patrick et al., 2016). We suggest that fist-RNA interactions contribute to this upstream sensing mechanism.

Coupled with the fist-RNA interactions, we detect a conformational change in the arm of the arch. Indeed, fist-RNA interactions drive this conformational change (**Figure 6**). Although the precise nature of the rearrangement of the arm is unclear, the result is that the fist and recA2 come together to place Mtr4 in a ‘closed-arch’ conformation. This contrasts the dramatically open conformation of the arch in Mtr4-exosome complexes (Schuller et al., 2018; E.-M. Weick et al., 2018). While the relationship between arch conformation and Mtr4 function are not completely understood, it is worth reiterating that we are monitoring initial RNA recognition events while the exosome-bound structures represent later states following translocation and delivery of the RNA to the exosome.

It seems likely that arch function in Mtr4 RNA binding and unwinding is regulated by Mtr4 interacting partners in complexes such as TRAMP. Whether and how these complexes exploit any of the intrinsic substrate selectivity mechanisms of Mtr4 remains to be seen. We note that Mtr4 recruitment by ribosome processing factors is mediated through protein-protein interactions in the fist via the arch interacting motifs of Nop53, Utp18 and NVL (Du et al., 2020; Lingaraju et al., 2019). The N-terminus of Air2 (a component of TRAMP) also binds to this same region (Falk et al., 2014). Future studies are needed to better understand the interplay between RNA, Mtr4 and Mtr4-interacting proteins. The application of HDX offers a promising path forward to characterization of this dynamic system.

## Materials and Methods

### Protein Preparation

The *Saccharomyces cerevisiae* Mtr4^ΔF^ was constructed by removing residues 667-813 and inserting a three-glycine linker using the Q5 Site-Directed Mutagenesis Kit (NEB). All Mtr4 proteins were recombinantly expressed in *Escherichia coli* BL21-CodonPlus (DE3)-RIPL (Stratagene) using an autoinduction protocol (Studier, 2005). The purification of Mtr4 was carried out as previously described with the addition of a HiTrap DEAE Sepharose FF (GE) before gel filtration (Jackson et al., 2010). RNase contamination was monitored with an RNaseAlert® kit (IDT) following heparin chromatography and at each subsequent chromatography step. Only RNase-free fractions were retained. Purified Mtr4 with 30% (v/v) glycerol was flash frozen in liquid nitrogen for storage.

### RNA Preparation

tRNA_i_^Met^ is similar to the native tRNA_i_^Met^ sequence from *S. cerevisiae* with a CCA cap and poly(A) sequence on the 3’ end. The sequence also includes non-native 5’ and 3’ cloning artifacts. The tRNA_i_^Met^ was generated using in vitro transcription with T7 RNA polymerase (ThermoFisher Scientific) on a plasmid linearized after the tRNA_i_^Met^ sequence. The tRNA_i_^Met^ was purified under non-denaturing conditions as previously described (Easton et al., 2010). Briefly, the transcription reaction was loaded onto HiTrap DEAE Sepharose FF (GE) and the tRNA_i_^Met^ eluted off using a NaCl gradient. All other RNAs were synthesized commercially (Integrated DNA Technologies, Inc.).

### Hydrogen-Deuterium Exchange Mass Spectrometry

Samples for the initial experiment (**Figure 1-2**) were prepared from stocks of 12.5 μM Mtr4^WT^ with or without 67.5 μM tRNA_i_^Met^ (5.4-fold). Samples for subsequent experiments (**Figure 3-6**) were prepared from stocks of 12.5 μM Mtr4^WT^ or Mtr4^ΔF^ with or without 25 μM RNA (2-fold). In both cases, stocks were prepared in 50 mM HEPES, 100 mM NaCl, 0.5 mM MgCl_2_, 0.2 mM TCEP, pH 6.5 and incubated for 10 min at 25°C. Following incubation, samples were diluted 2:23 with the same buffer containing H_2_O for controls (pH_read_ = 6.5 at 25°C) or D_2_O for exchange samples (pH_read_ = 6.1 at 25 °C; final D_2_O of 92%). Exchange proceeded at 25°C for 10 s, 10^2^ s, 10^3^ s, 10^4^ s, or 10^5^ s. Exchange was quenched by mixing samples 1:1 with cooled 1% (v/v) formic acid, 3.84 M guanidinium chloride, pH 1.75 (1:1 mix had a final pH of 2.3 at 0°C) and flash frozen in liquid nitrogen. Samples were prepared in triplicate and stored at −80°C.

Samples were thawed for 50 s immediately prior to injection into a Waters™ HDX manager in line with a SYNAPT G2-Si. In the HDX manager, samples were digested by *Sus scrofa* Pepsin A (Waters™ Enzymate BEH) at 15 °C and the peptides trapped on a C4 pre-column (Waters™ Acquity UPLC Protein BEH C4) at 1°C using a flowrate of 100 μL/min for 3 min. The chromatography buffer was 0.1 % (v/v) formic acid. Peptides were then separated over a C18 column (Waters™ Acquity UPLC BEH) at 1°C and eluted with a linear 3-40% (v/v) acetonitrile gradient using a flowrate of 40 uL/min for 7 min. Samples were injected in a random order.

Mass spectrometry data were acquired using positive ion mode in either HDMS or HDMS^E^ mode. Peptide identification of water-only control samples was performed using data-independent acquisition in HDMS^E^ mode. Peptide precursor and fragment data were collected via collision-induced dissociation at low (6V) and high (ramping 22-44 V) energy. HDMS mode was used to collect low energy ion data for all deuterated samples. All samples were acquired in resolution mode. Capillary voltage was set to 2.8 kV for the sample sprayer. Desolvation gas was set to 650 L/hour at 175 °C. The source temperature was set to 80°C. Cone and nebulizer gas was flowed at 90 L/hour and 6.5 bar, respectively. The sampling cone and source offset were both set to 30 V.Data were acquired at a scan time of 0.4 s with a range of 100-2000 m/z. Mass correction was done using [Glu1]-fibrinopeptide B as a reference. For ion mobility, the wave velocity was 700 ms^−1^ and the wave height was 40 V.

Raw data of Mtr4 water-only controls were processed by PLGS (Waters™ Protein Lynx Global Server 3.0.2) using a database containing *S. scrofa* Pepsin A and *S. cerevisiae* Mtr4^WT^ or Mtr4^ΔF^. In PLGS, the minimum fragment ion matches per peptide was 3 and methionine oxidation was allowed. The low and elevated energy thresholds were 250 and 50 counts respectively, and overall intensity threshold was 750 counts. DynamX 3.0 was used to search the deuterated samples for peptides with 0.3 products per amino acid and 1 consecutive product found in 2 out of 7-10 controls. Data were manually refined. Structural images were made using scripts from DynamX in Pymol (Schrödinger, LLC, 2015), and heat maps were made using HD-eXplosion (Zhang et al., 2021). To allow access to the HDX data of this study, the HDX data summary table (Table S1) and the HDX data table (Table S2) are included.

### Affinity Measurements

Dissociation constants (K_d_) between Mtr4 and RNA were measured using an electrophoretic mobility shift assay. Various concentrations of Mtr4^WT^ or Mtr4^ΔF^ were mixed with 60 nM RNA in binding buffer (40 mM MOPS, 100 mM NaCl, 2.5 mM MgCl_2_, 5% (v/v) glycerol, 0.01% (v/v) nonidet-P40 substitute, 1 U/μl of Ribolock (ThermoFisher Scientific), 2 mM dithiothreitol, pH 6.5). Samples were incubated on ice for 45-60 min to reach equilibrium and diluted 1:4 with loading buffer (20% (v/v) glycerol, 0.1% (w/v) bromophenol blue, 0.1% (v/v) xylene cyanol). Aliquots were separated on a native 4-20% gradient polyacrylamide TBE gel for 80-210 min at 120 V. The gel was stained with SYBR Gold Nucleic Acid Gel stain (Invitrogen) and imaged with a ChemiDoc MP Imaging system (Bio-Rad Laboratories). The fraction of RNA bound was quantified by densitometry using Image Lab (Bio-Rad Laboratories) and fit to a quadratic 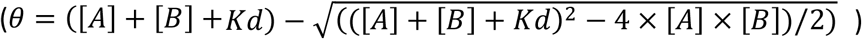, where *θ*is fraction bound, [A] is concentration of Mtr4, [B] is the concentration of RNA, and *Kd* is the dissociation constant. Data were fit using KaleidaGraph (Synergy Software).

### RNA Unwinding Assays

Pre-steady state unwinding assays were performed essentially as described (Jia et al., 2012; Morales et al., 2018; Taylor et al., 2014). Unwinding activity was determined by monitoring the displacement of a 16-nt RNA labeled with a fluorescein on its 5’ end, from a longer unlabeled RNA strand when incubated with Mtr4 at saturating levels of ATP. Unwinding reactions were performed in 40 mM MOPS, 100 mM NaCl, 0.5 mM MgCl_2_, 5% (v/v) glycerol, 0.01% (v/v) nonidet-P40 substitute, 1 U/μl of Ribolock (ThermoFisher Scientific), 2 mM dithiothreitol, pH 6.5 at 30°C. 50-1200 nM of Mtr4 was incubated with 10 nM fluorescein-labeled duplex RNA and 100 nM DNA-trap oligonucleotide for 5 min. Reactions were initiated by the addition of 1.6 mM ATP with MgCl_2_. At various times aliquots were quenched with a 1:1 dilution in 1% (v/v) SDS, 5 mM EDTA, 20% (v/v) glycerol. Samples were separated on a 15% native polyacrylamide TTE gel for 80 min at 120 V. Gels were imaged with a ChemiDoc MP Imaging system (Bio-Rad Laboratories).

## Supporting information

Supplemental Images and Tables

## Acknowledgements

This work was supported by start-up funds from The University of Texas at Dallas (S.D.) and a R01 from the National Institute of General Medical Sciences, NIH (R01GM117311 to S.J.). We thank members of the D’Arcy and Johnson Groups for critical discussions.

