## Supplemental Images and Tables for "Hydrogen-deuterium exchange mass spectrometry of Mtr4 with diverse RNAs reveals substrate-dependent dynamics and interfaces in the arch"

**Supplementary Tables and Figures**

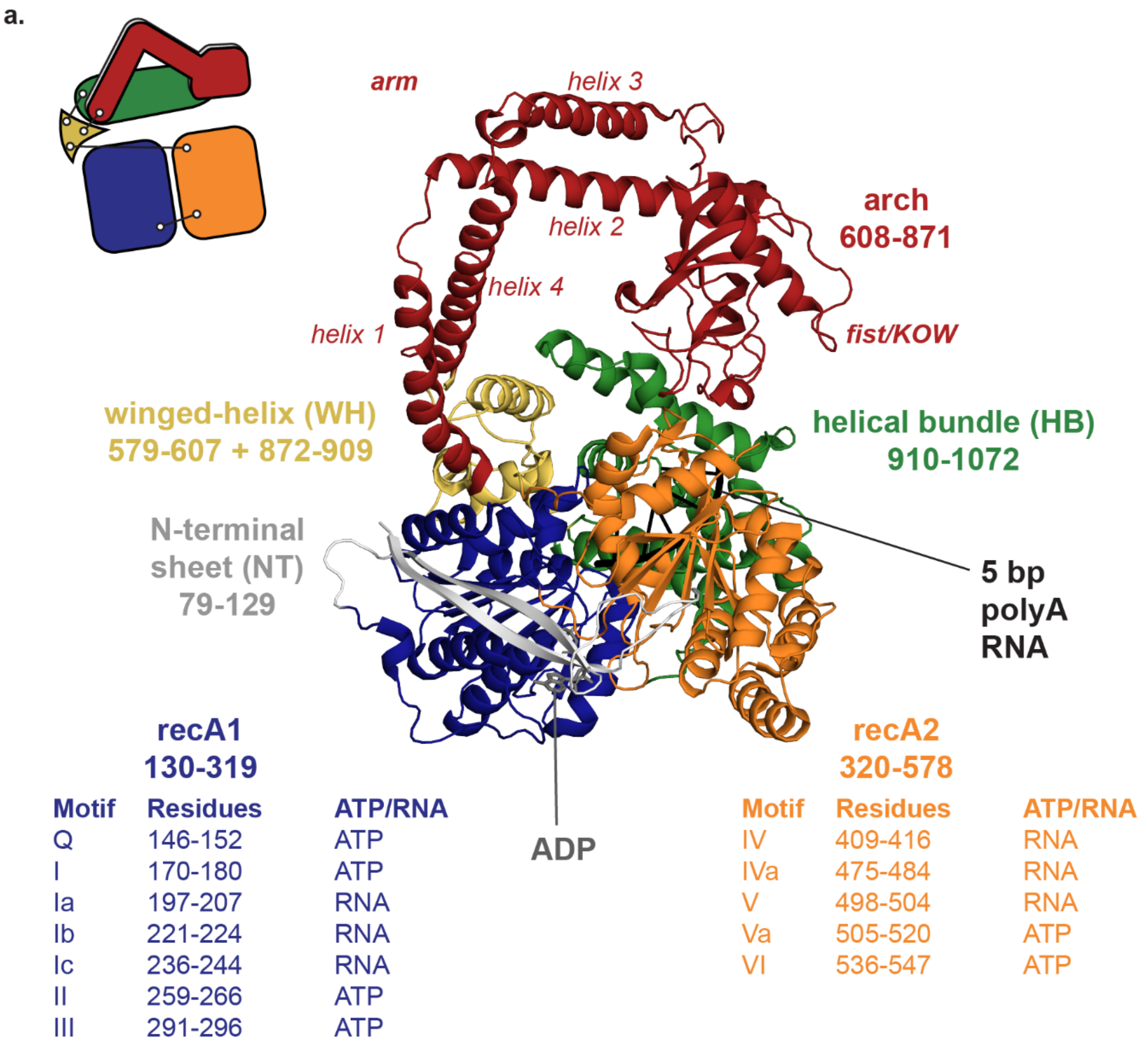

**b.**

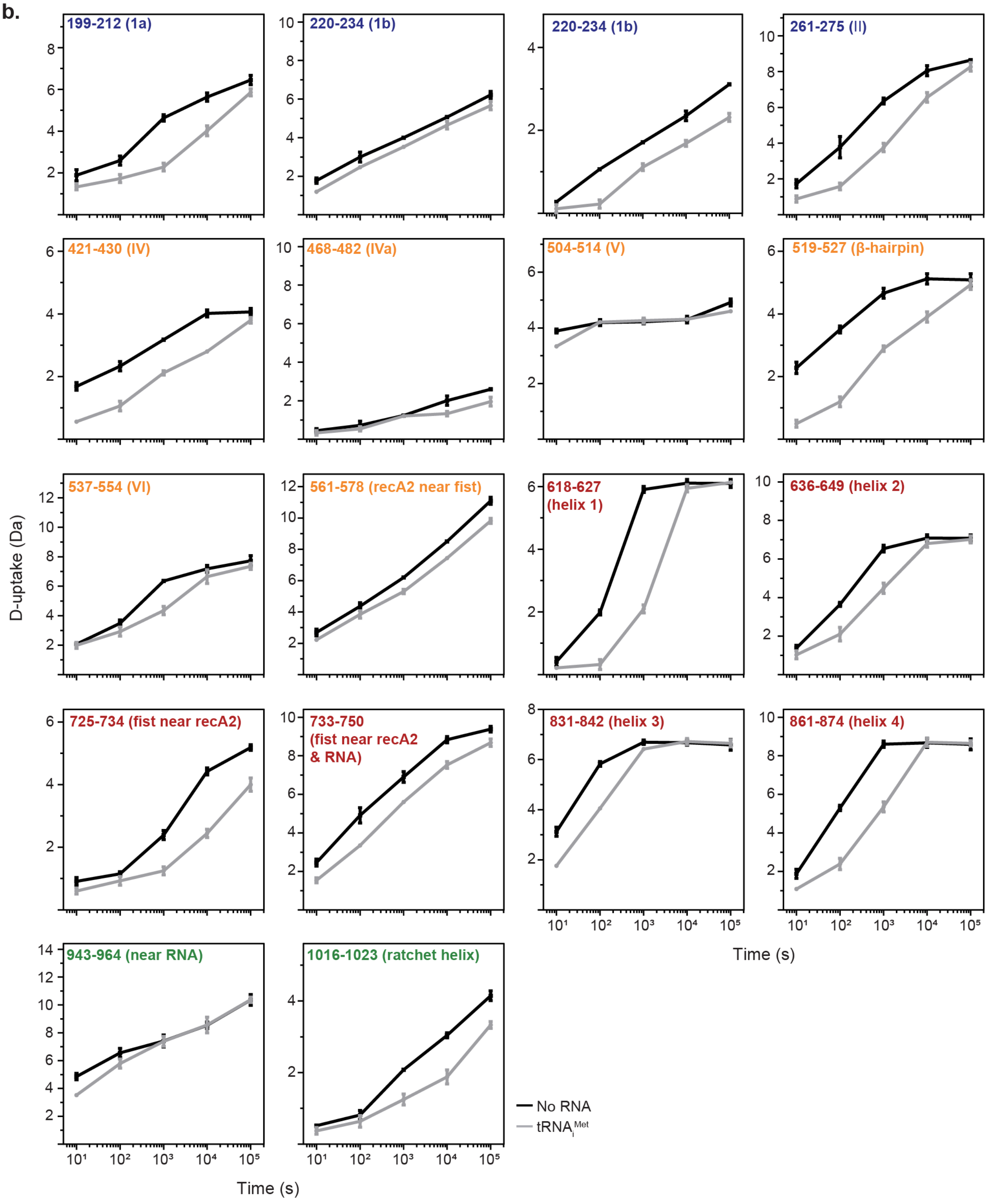
199-212 (1a)
220-234 (1b)
220-234 (1b)
261-275 (II)
421-430 (IV)
468-482 (IVa)
504-514 (V)
519-527 (β-hairpin)
537-554 (VI)
561-578 (recA2 near fist)
618-627 (helix 1)
636-649 (helix 2)
725-734 (fist near recA2)
733-750 (fist near recA2 & RNA)
831-842 (helix 3)
861-874 (helix 4)
943-964 (near RNA)
1016-1023 (ratchet helix)
D-uptake (Da)
Time (s)
No RNA
tRNAiMet

c.

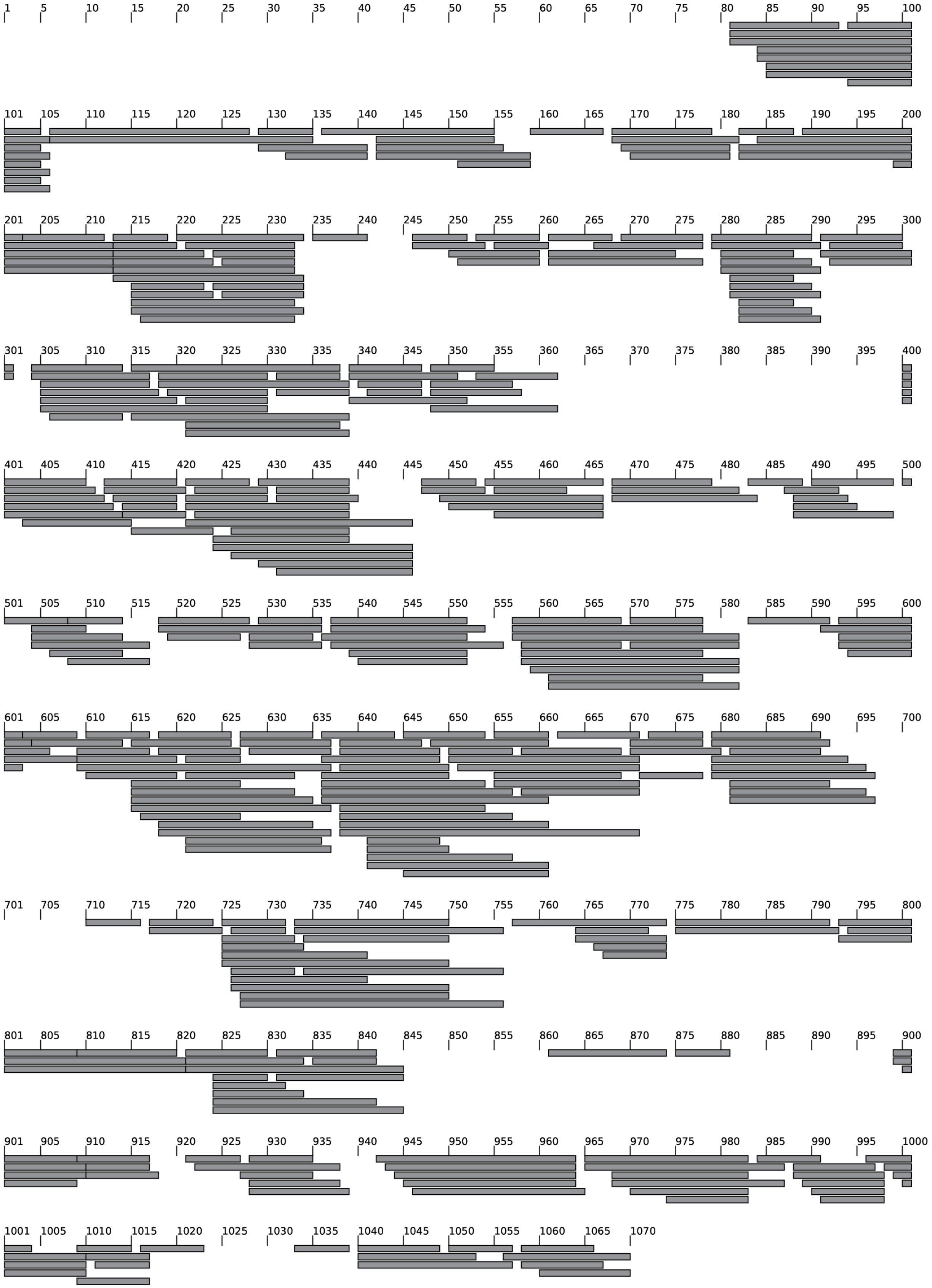
1 5 10 15 20 25 30 35 40 45 50 55 60 65 70 75 80 85 90 95 100
101 105 110 115 120 125 130 135 140 145 150 155 160 165 170 175 180 185 190 195 200
201 205 210 215 220 225 230 235 240 245 250 255 260 265 270 275 280 285 290 295 300
301 305 310 315 320 325 330 335 340 345 350 355 360 365 370 375 380 385 390 395 400
401 405 410 415 420 425 430 435 440 445 450 455 460 465 470 475 480 485 490 495 500
501 505 510 515 520 525 530 535 540 545 550 555 560 565 570 575 580 585 590 595 600
601 605 610 615 620 625 630 635 640 645 650 655 660 665 670 675 680 685 690 695 700
701 705 710 715 720 725 730 735 740 745 750 755 760 765 770 775 780 785 790 795 800
801 805 810 815 820 825 830 835 840 845 850 855 860 865 870 875 880 885 890 895 900
901 905 910 915 920 925 930 935 940 945 950 955 960 965 970 975 980 985 990 995 1000
1001 1005 1010 1015 1020 1025 1030 1035 1040 1045 1050 1055 1060 1065 1070

4

**Figure S1. Uptake plots and a peptide coverage map for Mtr4$^{WT}$ with 5.4-fold tRNA$_i^{Met}$. (a)** Mtr4 (PDB: 2XGJ) colored by domain with residue boundaries listed. Domains include N-terminal sheet (NT) (gray), recA1 (blue), recA2 (orange), winged-helix (WH) (yellow), arch (composed of arm and fist) (red), and helical bundle (green). ADP is gray sticks. Motifs for RNA binding or ATP binding and hydrolysis, and their residues in *S. cerevisiae* are listed. **(b)** Example uptake plots showing Mtr4$^{WT}$ alone (black) and with 5.4-fold tRNA$_i^{Met}$ (gray). Plotted uptake is the average of three replicates and error bars are plus or minus two standard deviations. The y-axis range is 80% of theoretical maximum uptake, assuming the N-terminal residue undergoes complete back-exchange. Data have not been corrected for back-exchange. **(c)** Peptides recovered from Mtr4$^{WT}$. The complete data set is shown in **Figure 1**. In this study, Mtr4 numbering does not include the start methionine (**Table S2**).

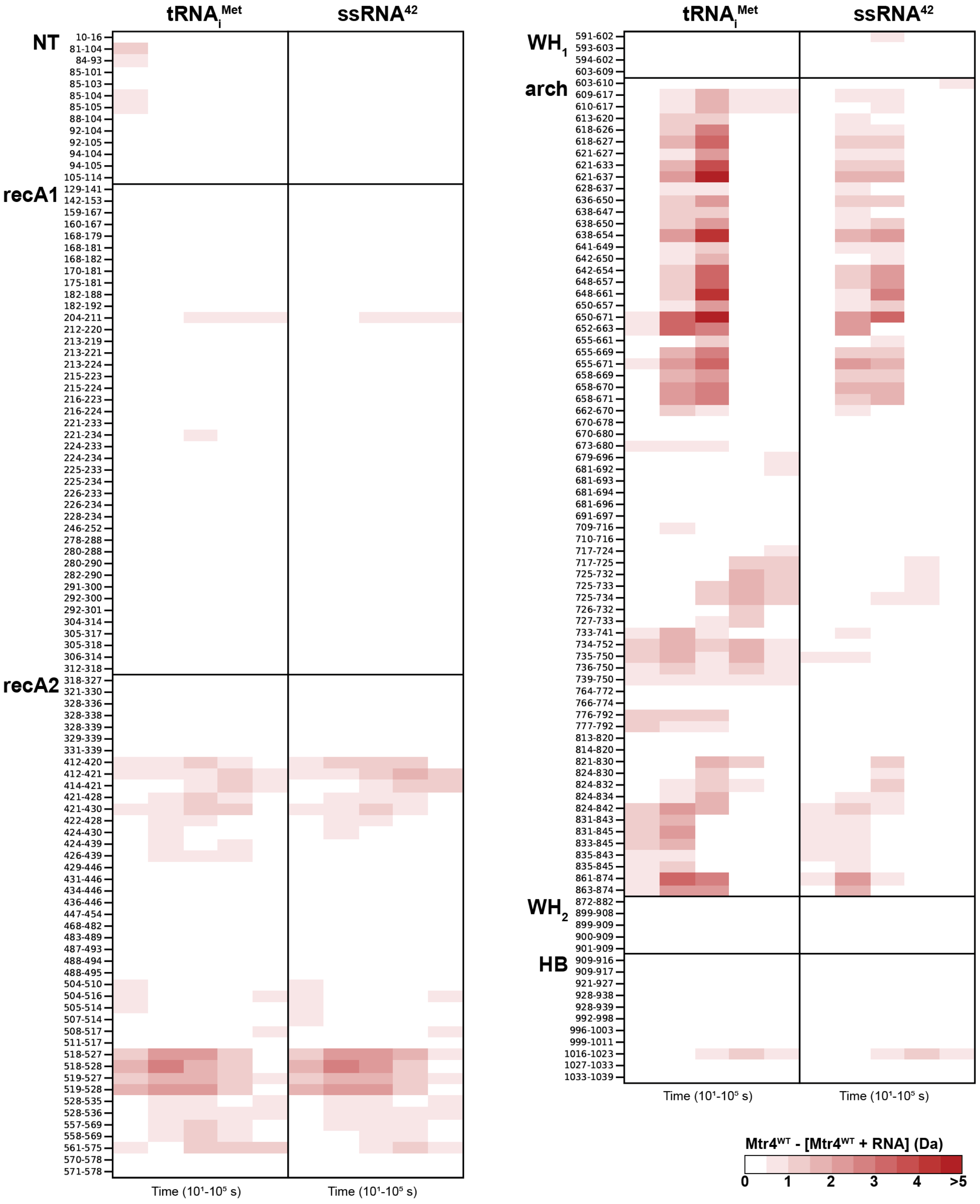
tRNAi^Met
ssRNA^42
NT
10-16
81-104
84-93
85-101
85-103
85-104
85-105
88-104
92-104
92-105
94-104
94-105
105-114
recA1
129-141
142-153
159-167
160-167
168-179
168-181
168-182
170-181
175-181
182-188
182-192
204-211
212-220
213-219
213-221
213-224
215-223
215-224
216-223
216-224
221-233
221-234
224-233
224-234
225-233
225-234
226-233
226-234
228-234
246-252
278-288
280-288
280-290
282-290
291-300
292-300
292-301
304-314
305-317
305-318
306-314
312-318
recA2
318-327
321-330
328-336
328-338
328-339
329-339
331-339
412-420
412-421
414-421
421-428
421-430
422-428
424-430
424-439
426-439
429-446
431-446
434-446
436-446
447-454
468-482
483-489
487-493
488-494
488-495
504-510
504-516
505-514
507-514
508-517
511-517
518-527
518-528
519-527
519-528
528-535
528-536
557-569
558-569
561-575
570-578
571-578
Time (10^1-10^5 s)
Time (10^1-10^5 s)
tRNAi^Met
ssRNA^42
WH1
591-602
593-603
594-602
603-609
arch
603-610
609-617
610-617
613-620
618-626
618-627
621-627
621-633
621-637
628-637
636-650
638-647
638-650
638-654
641-649
642-650
642-654
648-657
648-661
650-657
650-671
652-663
655-661
655-669
655-671
658-669
658-670
658-671
662-670
670-678
670-680
673-680
679-696
681-692
681-693
681-694
681-696
691-697
709-716
710-716
717-724
717-725
725-732
725-733
725-734
726-732
727-733
733-741
734-752
735-750
736-750
739-750
764-772
766-774
776-792
777-792
813-820
814-820
821-830
824-830
824-832
824-834
824-842
831-843
831-845
833-845
835-843
835-845
861-874
863-874
WH2
872-882
899-908
899-909
900-909
901-909
HB
909-916
909-917
921-927
928-938
928-939
992-998
996-1003
999-1011
1016-1023
1027-1033
1033-1039
Time (10^1-10^5 s)
Time (10^1-10^5 s)
Mtr4^WT - [Mtr4^WT + RNA] (Da)
0
1
2
3
4
>5

**Figure S2. Protection in the arch of Mtr4$^{WT}$ was reduced for ssRNA$^{42}$ compared to tRNA$_i^{Met}$.** Heatmaps showing the difference in deuterium uptake between Mtr4$^{WT}$ alone and Mtr4$^{WT}$ with 2-fold tRNA$_i^{Met}$ or ssRNA$^{42}$ for all recovered peptides. The time points of exchange (x-axis) were 10 s, $10^2$ s, $10^3$ s, $10^4$ s, and $10^5$ s. Red blocks indicate a difference ≥0.5 Da with a p-value ≤0.01 in a Welch's t-test (n=3). Figure was created using HD-eXplosion (Zhang et al., 2021). Part of the data set is in **Figure 3**.

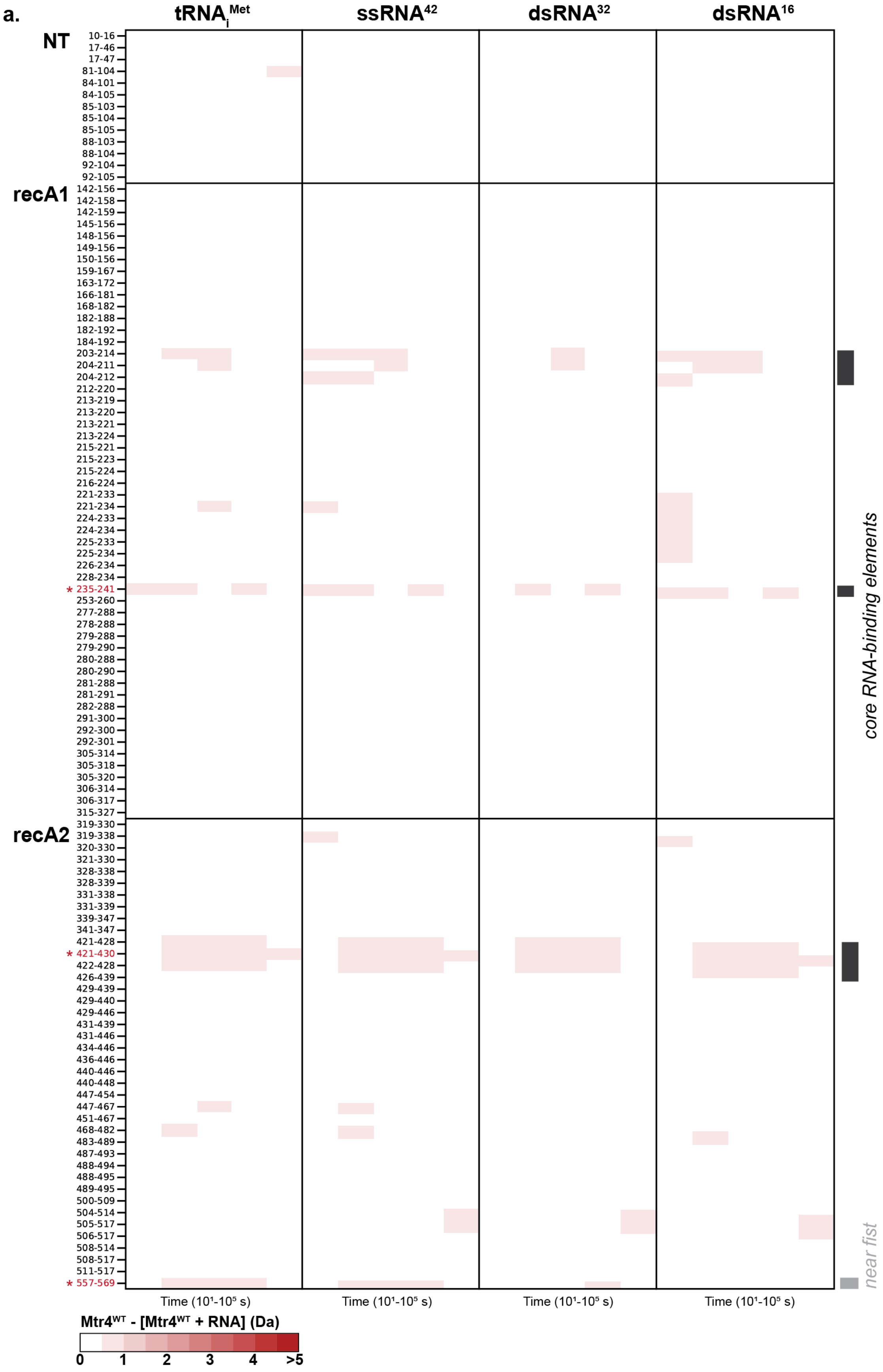
a.
tRNAiMet
ssRNA42
dsRNA32
dsRNA16
NT
10-16
17-46
17-47
81-104
84-101
84-105
85-103
85-104
85-105
88-103
88-104
92-104
92-105
recA1
142-156
142-158
142-159
145-156
148-156
149-156
150-156
159-167
163-172
166-181
168-182
182-188
182-192
184-192
203-214
204-211
204-212
212-220
213-219
213-220
213-221
213-224
215-221
215-223
215-224
216-224
221-233
221-234
224-233
224-234
225-233
225-234
226-234
228-234
* 235-241
253-260
277-288
278-288
279-288
279-290
280-288
280-290
281-288
281-291
282-288
291-300
292-300
292-301
305-314
305-318
305-320
306-314
306-317
315-327
recA2
319-330
319-338
320-330
321-330
328-338
328-339
331-338
331-339
339-347
341-347
421-428
* 421-430
422-428
426-439
429-439
429-440
429-446
431-439
431-446
434-446
436-446
440-446
440-448
447-454
447-467
451-467
468-482
483-489
487-493
488-494
488-495
489-495
500-509
504-514
505-517
506-517
508-514
508-517
511-517
* 557-569
core RNA-binding elements
near fist
Time (10¹-10⁵ s)
Time (10¹-10⁵ s)
Time (10¹-10⁵ s)
Time (10¹-10⁵ s)
Mtr4WT - [Mtr4WT + RNA] (Da)
0
1
2
3
4
>5

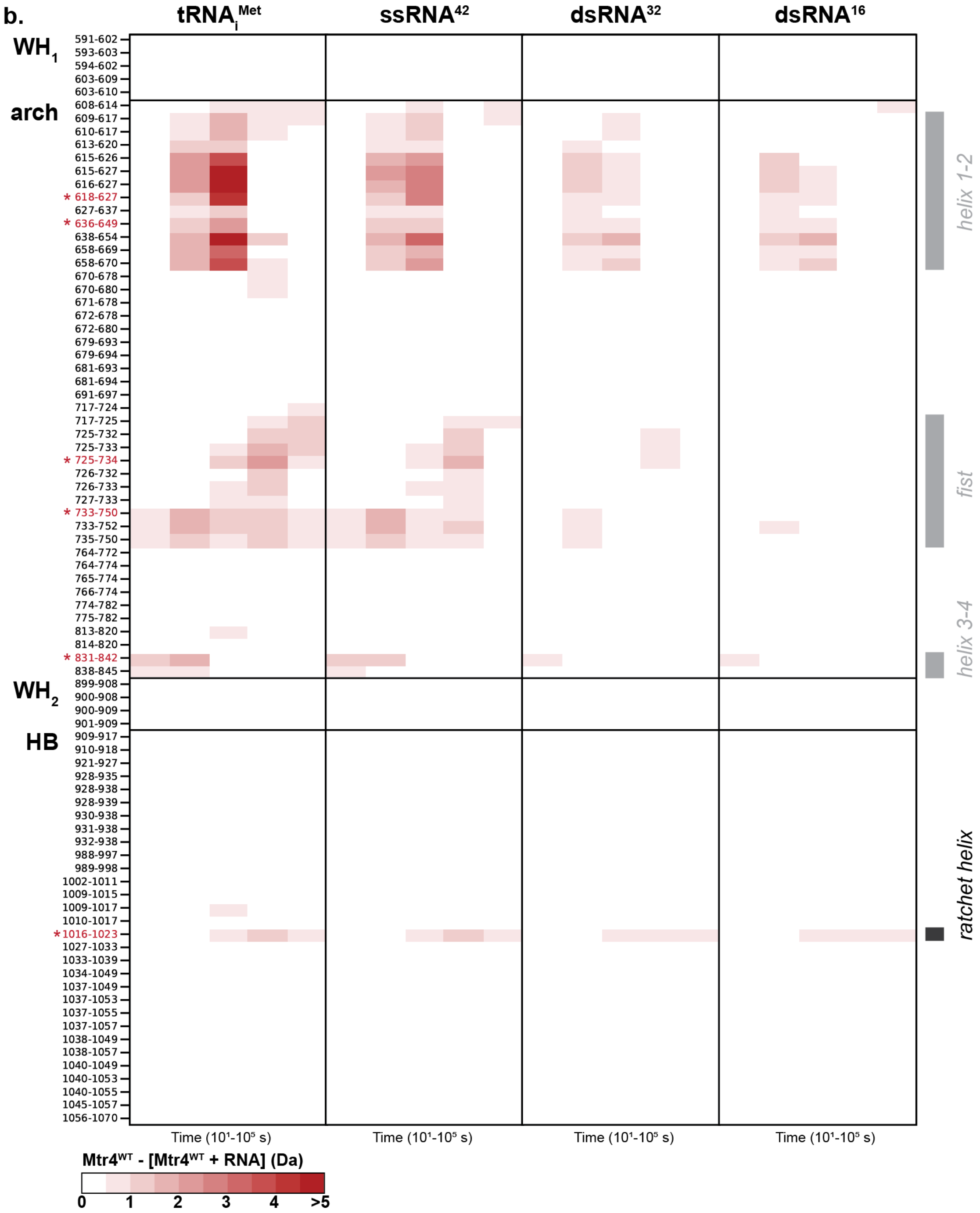

b.
tRNAiMet
ssRNA42
dsRNA32
dsRNA16
WH1
arch
WH2
HB
591-602
593-603
594-602
603-609
603-610
608-614
609-617
610-617
613-620
615-626
615-627
616-627
* 618-627
627-637
* 636-649
638-654
658-669
658-670
670-678
670-680
671-678
672-678
672-680
679-693
679-694
681-693
681-694
691-697
717-724
717-725
725-732
725-733
* 725-734
726-732
726-733
727-733
* 733-750
733-752
735-750
764-772
764-774
765-774
766-774
774-782
775-782
813-820
814-820
* 831-842
838-845
899-908
900-908
900-909
901-909
909-917
910-918
921-927
928-935
928-938
928-939
930-938
931-938
932-938
988-997
989-998
1002-1011
1009-1015
1009-1017
1010-1017
* 1016-1023
1027-1033
1033-1039
1034-1049
1037-1049
1037-1053
1037-1055
1037-1057
1038-1049
1038-1057
1040-1049
1040-1053
1040-1055
1045-1057
1056-1070
helix 1-2
fist
helix 3-4
ratchet helix
Time (10¹-10⁵ s)
Time (10¹-10⁵ s)
Time (10¹-10⁵ s)
Time (10¹-10⁵ s)
Mtr4WT - [Mtr4WT + RNA] (Da)
0
1
2
3
4
>5

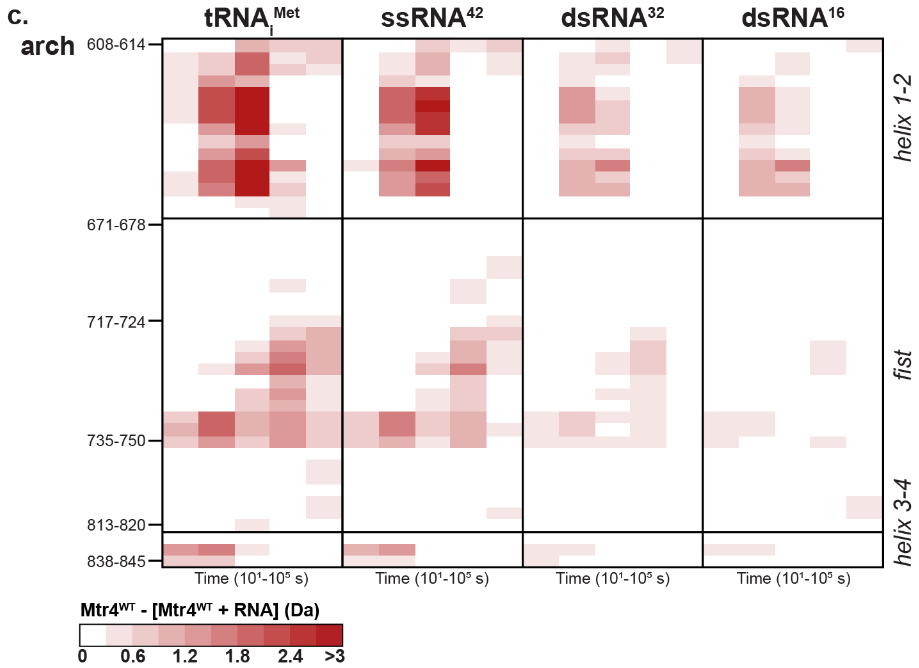


**Figure S3. The degree of protection in the arch but not the helicase core of $Mtr4^{WT}$ varied with RNA type.** Heatmaps showing the difference in deuterium uptake between $Mtr4^{WT}$ alone and $Mtr4^{WT}$ with 2-fold $tRNA_i^{Met}$, $ssRNA^{42}$, $dsRNA^{32}$, or $dsRNA^{16}$, for all recovered peptides from **(a)** NT, recA1 and recA2; and **(b)** WH, arch, and HB. The time points of exchange (x-axis) were 10 s, $10^2$ s, $10^3$ s, $10^4$ s, and $10^5$ s. Red blocks indicate a difference ≥0.5 Da with a p-value ≤0.01 in a Welch's t-test (n=3). Figure was created using HD-eXplosion (Zhang et al., 2021). Peptides with uptake plots shown in **Figure 4** are indicated (red and *) and Mtr4 features in the helicase core (black) or arch (gray) are labeled. Peptides from the arch are shown with a 0.5 Da cutoff (**Figure 5**), as well as with a 0.3 Da cutoff **(c)**.

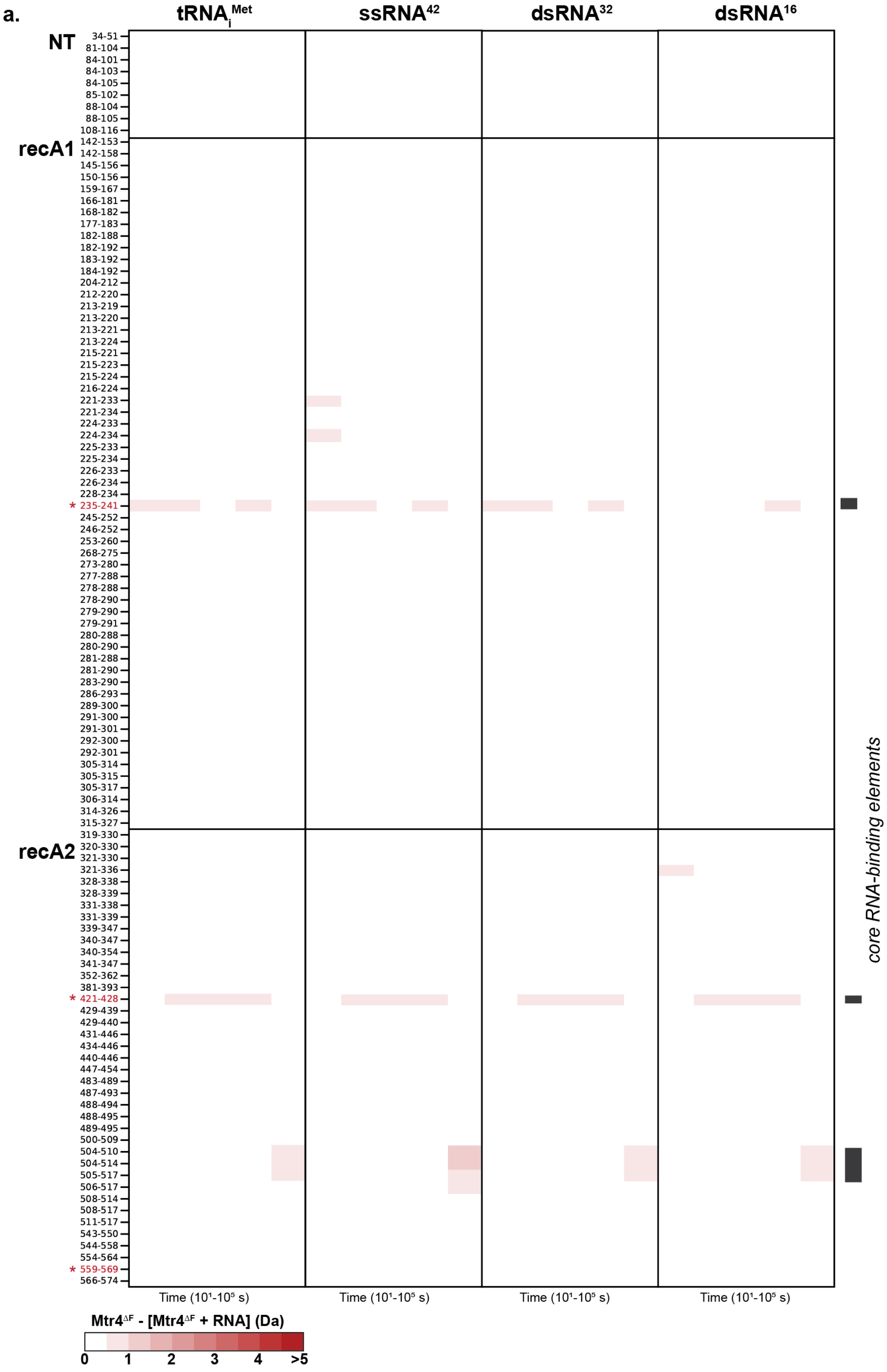
a.
tRNAiMet
ssRNA42
dsRNA32
dsRNA16
NT
34-51
81-104
84-101
84-103
84-105
85-102
88-104
88-105
108-116
recA1
142-153
142-158
145-156
150-156
159-167
166-181
168-182
177-183
182-188
182-192
183-192
184-192
204-212
212-220
213-219
213-220
213-221
213-224
215-221
215-223
215-224
216-224
221-233
221-234
224-233
224-234
225-233
225-234
226-233
226-234
228-234
* 235-241
245-252
246-252
253-260
268-275
273-280
277-288
278-288
278-290
279-290
279-291
280-288
280-290
281-288
281-290
283-290
286-293
289-300
291-300
291-301
292-300
292-301
305-314
305-315
305-317
306-314
314-326
315-327
recA2
319-330
320-330
321-330
321-336
328-338
328-339
331-338
331-339
339-347
340-347
340-354
341-347
352-362
381-393
* 421-428
429-439
429-440
431-446
434-446
440-446
447-454
483-489
487-493
488-494
488-495
489-495
500-509
504-510
504-514
505-517
506-517
508-514
508-517
511-517
543-550
544-558
554-564
* 559-569
566-574
Time (10¹-10⁵ s)
Time (10¹-10⁵ s)
Time (10¹-10⁵ s)
Time (10¹-10⁵ s)
core RNA-binding elements
Mtr4ΔF - [Mtr4ΔF + RNA] (Da)
0
1
2
3
4
>5

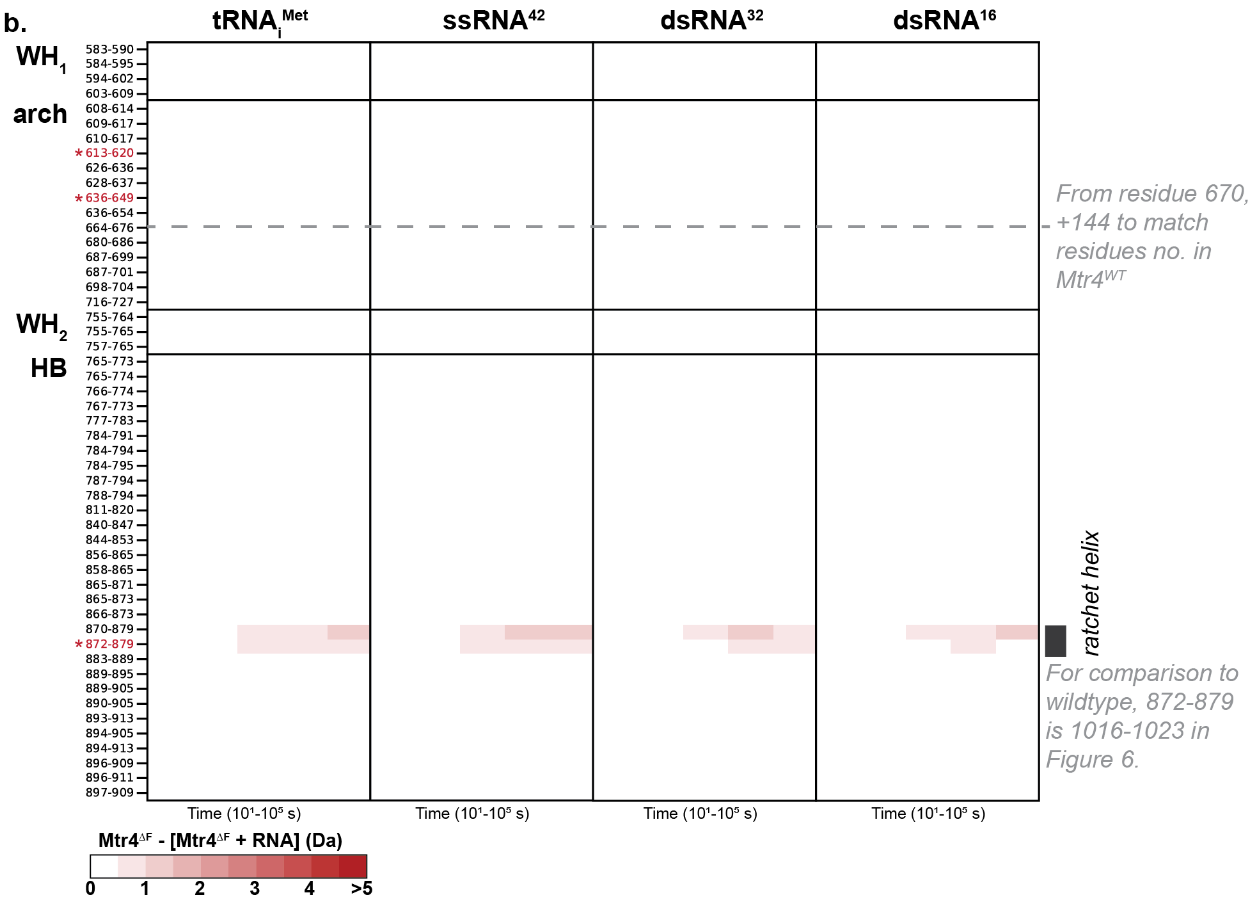


**Figure S4. Deletion of the fist in Mtr4$^{\Delta F}$ caused a loss of protection in the arch but not the helicase core.** Heatmaps showing the difference in deuterium uptake between Mtr4$^{\Delta F}$ alone and Mtr4$^{\Delta F}$ with 2-fold tRNA$_i^{Met}$, ssRNA$^{42}$, dsRNA$^{32}$, or dsRNA$^{16}$, for all recovered peptides from **(a)** NT, recA1 and recA2; and **(b)** WH, arch, and HB. The time points of exchange (x-axis) were 10 s, $10^2$ s, $10^3$ s, $10^4$ s, and $10^5$ s. Red blocks indicate a difference ≥0.5 Da with a p-value ≤0.01 in a Welch's t-test (n=3). Figure was created using HD-eXplosion (Zhang et al., 2021). Peptides with uptake plots shown in **Figure 6** are indicated (red and *) and Mtr4 features in the helicase core (black) are labeled.

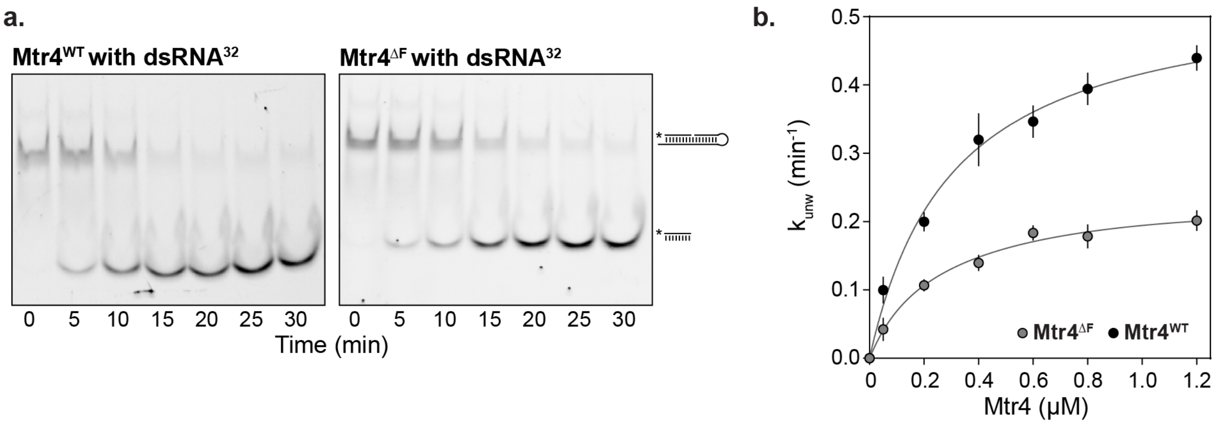


**Figure S5. Deletion of the fist in Mtr4$^{\Delta F}$ caused slower unwinding of dsRNA$^{32}$. (a)** Representative 15% native polyacrylamide TTE gels for unwinding of dsRNA$^{32}$ by Mtr4$^{WT}$ and Mtr4$^{\Delta F}$. Unwinding was monitored by quantifying the fluorescence signal of bands representing duplex and single-stranded nucleic acid. The displaced strand contains a 5′ fluorescein tag indicated by an *. **(b)** Rate constants ($k_{unw}$) for unwinding dsRNA$^{32}$ were plotted against Mtr4 concentration for Mtr4$^{WT}$ (black dots) and Mtr4$^{\Delta F}$ (gray dots). Rate constants were the average of at least three independent reactions and the error bars are plus or minus one standard deviation. Data were fit to a binding isotherm ($k_{unw} = k_{unw}^{max}$ [E]/([E] + $K_{1/2}^{max}$)) where [E] is Mtr4 concentration, $k_{unw}^{max}$ is the maximum unwinding rate, and $K_{1/2}^{max}$ is the functional affinity, as described in (Jia et al., 2012).

**Table S1. Summary of HDX Experiments**

| | **Mtr4$^{WT}$**<br>**Figure 1-2** | **Mtr4$^{WT}$**<br>**Figure 3** | **Mtr4$^{WT}$**<br>**Figure 4-5** | **Mtr4$^{ΔF}$**<br>**Figure 6** |
|---|---|---|---|---|
| **Samples** | apo<br>$tRNA_i^{Met}$ | apo<br>$tRNA_i^{Met}$<br>$ssRNA^{42}$ | apo<br>$tRNA_i^{Met}$<br>$ssRNA^{42}$<br>$dsRNA^{32}$<br>$dsRNA^{16}$ | apo<br>$tRNA_i^{Met}$<br>$ssRNA^{42}$<br>$dsRNA^{32}$<br>$dsRNA^{16}$ |
| **[Mtr4] (µM)** | 1 | 1 | 1 | 1 |
| **[RNA] (µM)** | 5.4 | 2 | 2 | 2 |
| **HDX buffer** | 50 mM HEPES, 100 mM NaCl, 0.5 mM $MgCl_2$, 0.2 mM TCEP, $pH_{read}$=6.1 | | | |
| **HDX time course** | $10^1$, $10^2$, $10^3$, $10^4$, $10^5$ s at 25°C | | | |
| **HDX control samples** | Mtr4$^{WT}$ | | | Mtr4$^{ΔF}$ |
| **Back exchange** | ~30% | | | |
| **Significance cutoffs** | ΔHDX > 0.5 Da and p-value < 0.01 in Welch's t-test (n=3) | | | |
| **# Peptides** | 339 | 188 | 191 | 161 |
| **Sequence coverage (%)** | 81 | 58 | 58 | 64 |
| **Average peptide length** | 12.6 | 9.86 | 11.08 | 9.7 |
| **Peptide redundancy** | 5.14 | 3.21 | 3.38 | 2.82 |
| **Technical replicates** | 3 | 3 | 3 | 3 |
| **Mean SD** | 0.064 | 0.061 | 0.051 | 0.050 |

**Table S2. Deuterium uptake data for all HDX experiments**

See attached excel file.

**Table S3. RNA Constructs**

Schematics and sequences of RNA constructs. The first and last base of the structured tRNA are indicated by subscripts. 3' overhangs are underlined. For dsRNA, complementary regions are colored gray.

| RNA Schematic | Sequence |
|---|---|
| **tRNA$_i^{Met}$** 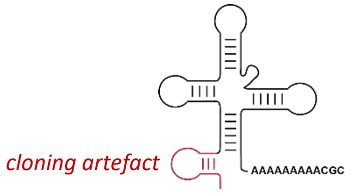  | 5'GGGGUUCACUGCCGUAUAGGCAGA$_1$GCGCCGUGGCGCAG UGGAAGCGCGCAGGGCUCAUAACCCUGAUGUCCUCGGAUC GAAACCGAGCGGCGCC$_{71}$AAAAAAAAACGC3' |
| **ssRNA[42]** 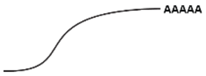  | 5'GCGUCUUUACGGUGCUUAAAACAAAACAAAACAAAACA AAAA3' |
| **dsRNA[32]** 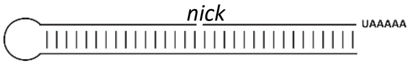  | 5' GCAGGAGCCUAGCUGAAAGAUGUCCUCGGAUCGACGCG UCUUUACGGUGCUUAAAAA3'<br>3'CGCAGAAAUGCCACGA5' |
| **dsRNA[16]** 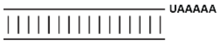  | 5'GCGUCUUUACGGUGCUUAAAAA3'<br>3'CGCAGAAAUGCCACGA5' |
